# RyR2-Mediated Ca^2+^ Leak Drives Progressive Arrhythmogenic Cardiomyopathy

**DOI:** 10.64898/2026.11.02.763829

**Authors:** Jingjing Zheng, Daniela Ponce-Balbuena, Sergio Estrada, Li Xiao, Erick B. Ríos Pérez, Achal M. Gowda, Claudia E. Korcarz, Kristin M. Hansen, Daniel J. Blackwell, Madelaine P. Thorpe, Wendell S. Akers, Jingshing Wu, Jeffrey N. Johnston, Ana M. Gómez, Jean-Pierre Benitah, Bjorn C. Knollmann, Y. Eugene Chen, Héctor H. Valdivia, Francisco J. Alvarado

**Author notes:** **Corresponding Author:** Francisco J. Alvarado, PharmD, PhD, 1111 Highland Ave, 8555 WIMR II, Madison, WI 53705.

## Abstract

**Background:** Abnormal sarcoplasmic reticulum (SR) Ca^2+^ release through ryanodine receptor 2 (RyR2) is a well-established mechanism of ventricular arrhythmias. Whether persistent RyR2 dysfunction can also initiate the structural remodeling characteristic of arrhythmogenic cardiomyopathy (ACM) remains unknown. We tested the hypothesis that chronic RyR2-mediated Ca^2+^ leak is sufficient to couple electrical instability to progressive cardiomyopathy.

**Methods:** We generated a rabbit carrying the severe gain-of-function RyR2-V2475F mutation and characterized disease progression using longitudinal cardiac phenotyping, continuous electrocardiographic monitoring, cellular electrophysiology and Ca^2+^ imaging, histopathology, biochemical analyses, and single-nucleus RNA sequencing (snRNA-seq). Therapeutic potential was evaluated by continuous treatment with the selective RyR2 inhibitor *ent*-verticilide for two months.

**Results:** RyR2-V2475F rabbits developed spontaneous ventricular tachyarrhythmias, sudden cardiac death, and progressive right ventricular (RV)-dominant fibrofatty cardiomyopathy recapitulating major features of human ACM. Malignant ventricular arrhythmias preceded overt structural remodeling and were associated at the cellular level with early spontaneous Ca^2+^ release and early afterdepolarizations. Early disease was characterized by remodeling of excitation-contraction coupling with preserved SR Ca^2+^ load and contractile function despite RyR2 dysfunction; upon disease progression, these compensatory changes were accompanied by SR Ca^2+^ depletion, impaired RV contractility and reduced left ventricular filling. Single-nucleus transcriptomics identified early fibroblast expansion and predicted cardiomyocyte-centered paracrine signaling before advanced fibrofatty infiltration. Cardiomyocytes also exhibited an altered lipid-metabolic transcriptome and intracellular lipid accumulation, accompanied by enhanced mitochondrial Ca^2+^ uptake. Selective RyR2 inhibition markedly reduced ventricular tachycardia burden and preserved RV contraction and LV filling during disease progression.

**Conclusions:** Severe RyR2 gain-of-function can lead to both malignant ventricular arrhythmias and progressive arrhythmogenic cardiomyopathy in a large-animal model, establishing persistent intracellular Ca^2+^ dysregulation as a mechanistic link between electrical and structural heart disease. The ability of selective RyR2 inhibition to suppress arrhythmias and preserve cardiac function identifies RyR2 as a potentially actionable therapeutic target in arrhythmogenic cardiomyopathy.

## INTRODUCTION

Arrhythmogenic cardiomyopathy (ACM) is a major cause of sudden cardiac death in young individuals. It is characterized by early-onset lethal ventricular arrhythmias followed by progressive cardiomyocyte loss, fibrofatty myocardial replacement and heart failure.^1^ Although inherited defects in desmosomal proteins account for most genetically defined cases, increasing evidence indicates that abnormal intracellular Ca^2+^ handling is a common feature across different subtypes of ACM.^1–5^ Enhanced sarcoplasmic reticulum (SR) Ca^2+^ leak through the cardiac ryanodine receptor (RyR2) has emerged as an early mechanism promoting ventricular arrhythmias, positioning RyR2 as a potential therapeutic target.^2,4,6^ However, it remains unresolved whether persistent RyR2 dysfunction contributes to the onset and progression of the structural disease in addition to triggering arrhythmias.

Evidence supporting a broader role for RyR2 in cardiomyopathy originates from both human genetics and experimental studies.^2,4,6–14^ Gain-of-function RyR2 mutations are classically associated with catecholaminergic polymorphic ventricular tachycardia (CPVT), an arrhythmia disorder that presents in the absence of overt structural remodeling.^15^ However, several studies have identified RyR2 variants in patients with different forms of inherited structural disease.^7–14^ Mouse models carrying relevant mutations have also been reported to develop structural abnormalities, including hypertrophic cardiomyopathy and left ventricular noncompaction, although these phenotypes are generally mild and the underlying mechanisms are not fully understood.^13,14^ Interestingly, mutations in RyR2 were linked to ACM even before a link with CPVT was established,^7,16^ but this association has been challenged in part because of the limited mechanistic support.^17^ Nonetheless, data suggest that RyR2 dysfunction may represent a nodal point where diverse genetic insults, including in RyR2 itself, converge to drive arrhythmia and structural remodeling of the heart. Yet, whether RyR2 dysfunction is sufficient to drive cardiomyopathy and ACM remains unconfirmed experimentally.

Efforts to answer this question have been limited, in part, by currently available animal models. In general, fibrofatty myocardial replacement, a hallmark feature of ACM, is difficult to recapitulate in mice.^1^ Mouse models carrying ACM-associated RyR2 mutations faithfully reproduce stress-induced ventricular arrhythmias but fail to develop the structural remodeling observed in patients.^17^ Furthermore, important differences in cardiac electrophysiology and Ca^2+^ handling between rodents and humans complicate mechanistic translation. Rabbit hearts exhibit action potential morphology, excitation-contraction coupling, and myocardial architecture, particularly epicardial fat, that more closely resemble those of humans (**Figure S1**), providing an advantageous platform for investigating RyR2-dependent disease mechanisms.

To test the hypothesis that RyR2 dysfunction can drive ACM in a model system physiologically closer to humans, we generated a rabbit harboring the severe gain-of-function mutation RyR2-V2475F.^18^ While heterozygotes have structurally normal hearts, homozygous animals show a cardiac phenotype consistent with human ACM. Disease progression involved structural remodeling that evolved from an early pre-overt stage with minimal remodeling to an advanced cardiomyopathy in addition to increased susceptibility to sudden cardiac death. Using extensive in vivo phenotyping, cellular electrophysiology, Ca^2+^ imaging, molecular analyses, and single-nucleus transcriptomics, we demonstrate a pattern of cardiac remodeling that became apparent during the early stage. It involved Ca^2+^ mishandling and arrhythmia, cardiomyocyte-specific metabolic remodeling and paracrine signaling affecting stromal cell populations, while traditional ACM pathways were unaffected. Furthermore, we show that long-term pharmacological RyR2 stabilization suppresses ventricular arrhythmias and preserves cardiac function in RyR2-V2475F rabbits. These data demonstrate, for the first time, that primary dysfunction in RyR2 caused by a point mutation can drive the phenotypic spectrum of ACM. Nonetheless, the role of RyR2 mutations in human ACM remains to be determined. The implications of our study span beyond this potentially rare link, positioning RyR2 dysfunction as a potential instigator of structural remodeling and establishing the channel as a promising therapeutic target for ACM and other cardiomyopathies.

## METHODS

RyR2-V2475F rabbits were generated via CRISPR/Cas9 at the Center for Advanced Models and Translational Sciences and Therapeutics (CAMTraST) at the University of Michigan. All animal procedures were approved by the IACUC of the University of Wisconsin School of Medicine and Public Health (M005944). Detailed methods can be found in the Supplementary Materials.

## RESULTS

### Chronic RyR2 dysfunction induces progressive arrhythmogenic cardiomyopathy

To test the hypothesis that RyR2-mediated SR Ca^2+^ leak is sufficient to drive structural cardiomyopathy in a species with human-like cardiac physiology, we generated rabbits harboring the severe gain-of-function RyR2-V2475F mutation (**Figure S2A**).^18^ In mice, homozygous inheritance is embryonic lethal.^18^ However, RyR2-V2475F rabbits reproduced in a Mendelian ratio (**Figure S2B**), indicating that the homozygous condition is compatible with embryonic development. Such species difference may result from the smaller contribution of SR Ca^2+^ to cardiac excitation-contraction coupling in larger mammals compared to rodents (**Figure S1C**).^19^ RyR2 channels isolated from homozygous rabbit hearts exhibited increased sensitivity to activation by Ca^2+^ (**Figure S3A-C**) consistent with previous data with recombinant protein.^18^ In contrast, RyR2 channels purified from heterozygous hearts showed similar Ca^2+^-dependent activation to WT controls (**Figure S3D-F**). This is likely a reflection of the heterogeneous population of RyR2 channels in which a tetramer can contain zero through four mutant subunits.^18^ Homozygous rabbits (VF), however, express a uniform population of gain-of-function RyR2 tetramers. This positions the homozygous condition as a robust proof-of-concept model for investigating the long-term consequences of chronic RyR2-mediated SR Ca^2+^ leak in vivo.

Notably, 32% (37/115) of the VF animals succumbed to sudden death (SD) at a median age of 17.7 weeks (**Figure S4**). Postmortem analysis revealed progressive myocardial degeneration characterized by cardiomyocyte loss, extensive fibrofatty replacement, and patchy inflammatory infiltrates (**Figure 1, Figure S5, Figure S6**). The pathology evolved from early right ventricular (RV) dilation without detectable fibrosis to RV-dominant fibrofatty infiltration, ultimately progressing to biventricular cardiomyopathy in older animals. Fibrosis and adipocyte accumulation were confirmed by Masson’s trichrome staining and anti-perilipin-1 immunohistochemistry, respectively (**Figure 1A-B**). Fibrofatty replacement originated in the epicardium and progressed toward the endocardium, a hallmark feature of ACM.^1^

**Figure 1.**
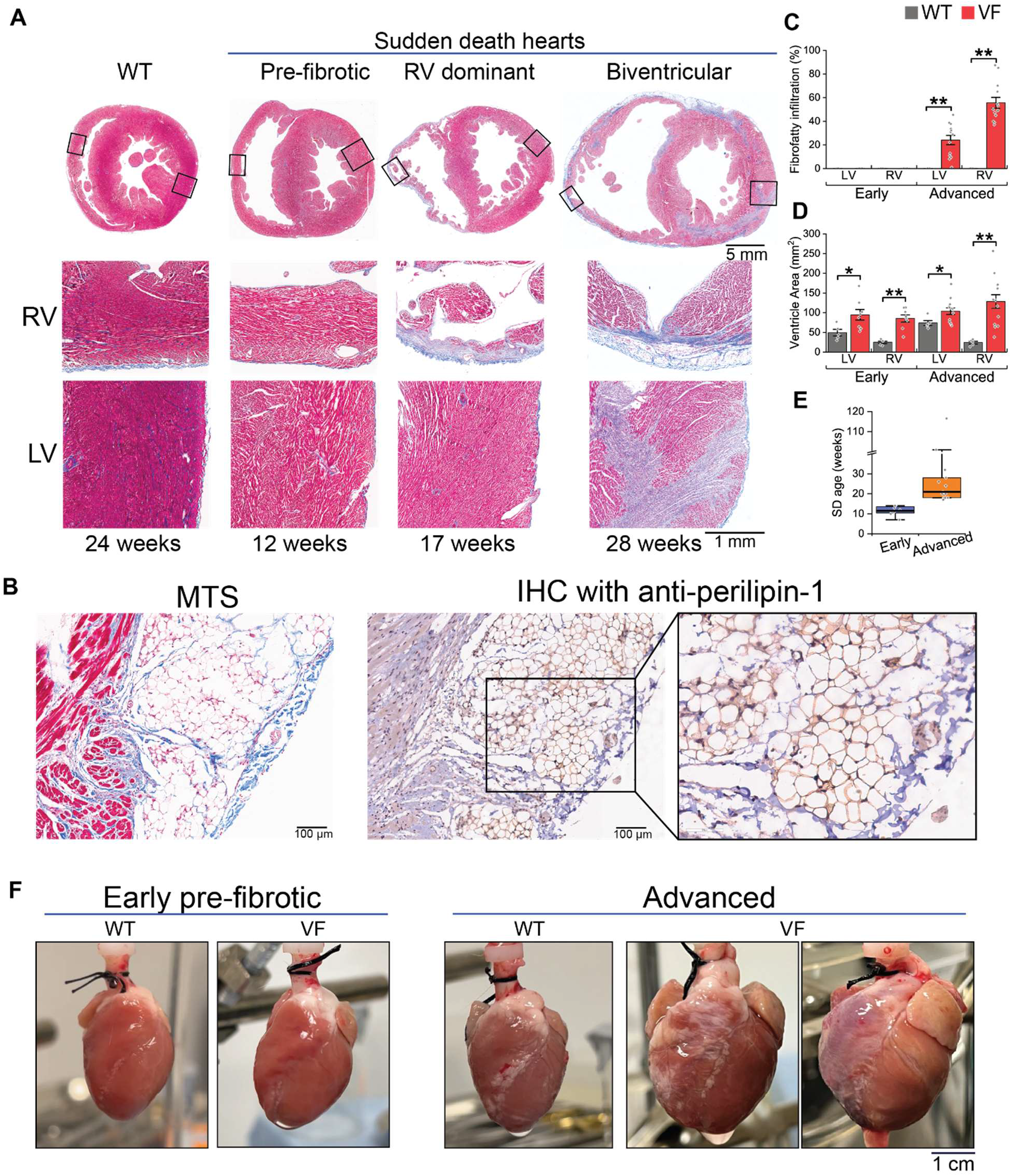
Sudden death and progressive right ventricular-dominant structural remodeling in RyR2-V2475F rabbits. **A**. Representative Masson’s trichrome staining (MTS) of heart sections from WT rabbits and RyR2-V2475F knock-in (VF) rabbits that experienced sudden death (SD). VF hearts exhibited progressive remodeling, ranging from mild RV dilation (early, pre-fibrotic stage) to RV-dominant wall thinning and fibrosis, and ultimately biventricular dilation with extensive fibrofatty infiltration (advanced stage). Myocardial loss followed a characteristic epicardium-to-endocardium pattern. Left: representative WT heart harvested at 24 weeks of age. Right: postmortem hearts from VF rabbits that experienced SD at the indicated ages. Top row: whole-heart transverse sections. Middle row: higher magnification of the RV (boxed region). Bottom row: higher magnification of the LV (boxed region). **B**. Histological validation of adipose infiltration in the LV of a VF rabbit at 28 weeks old. Masson’s Trichrome staining (MTS) and perilipin-1 immunohistochemistry (IHC) confirm fibrofatty replacement originating predominantly from the epicardium. **C-D**. Quantification of fibrofatty area (C) and ventricular perimeter (D) from tissue slides (mean±SEM). Mutant rabbits exhibit significant LV and RV dilation beginning at the early stage, with more pronounced enlargement in the RV. **E**. Age at sudden death in VF rabbits, stratified by early (N = 8) and advanced (N = 13) stages. **F**. Anterior view of Langendorff-perfused WT and VF hearts harvested from live animals at the early and advanced stages. (C-E: Early: WT = 5, VF = 8; Advanced: WT = 5, VF = 13. *P < 0.05, **P < 0.01; two-tailed t-test or Mann-Whitney test between stage and side matched WT and VF).

To define disease progression, we stratified the pathology into an early stage (before 14 weeks of age), featuring RV dilation without detectable fibrotic infiltration by histology, and an advanced stage (after 17 weeks of age) with extensive biventricular fibrofatty infiltrations (56±5% RV and 24±4% left ventricle [LV]) (**Figure 1C,D**). SD clustered between these stages, occurring at a median age of 11.5 weeks in the early stage and 21 weeks in the advanced stage (**Figure 1E**). Hearts collected from age-matched surviving VF cohorts recapitulated these stage-specific features, establishing a temporally resolved and experimentally traceable model of disease progression (**Figure 1F**).

### Electrical instability precedes overt fibrofatty remodeling and involves a distinct cellular mechanism

Continuous ECG monitoring in conscious animals revealed spontaneous ventricular arrhythmias in all VF rabbits independently of disease stage (**Figure 2A-B**), whereas no arrhythmias were observed in age-matched WT controls. The predominant arrhythmias included polymorphic ventricular tachycardia (Poly VT) and ventricular bigeminy, with occasional bidirectional and monomorphic VT (BDVT and Mono VT, respectively) (**Figure 2C**). In the case of a rabbit with SD, sustained VT degenerated into ventricular fibrillation followed by terminal asystole in the absence of overt fibrofatty infiltration (**Figure 2D**). Thus, RyR2-V2475F rabbits recapitulate the defining electrical and histopathological features of human ACM, including early ventricular arrhythmias followed by progressive epicardial-to-endocardial fibrofatty remodeling.^1^

**Figure 2.**
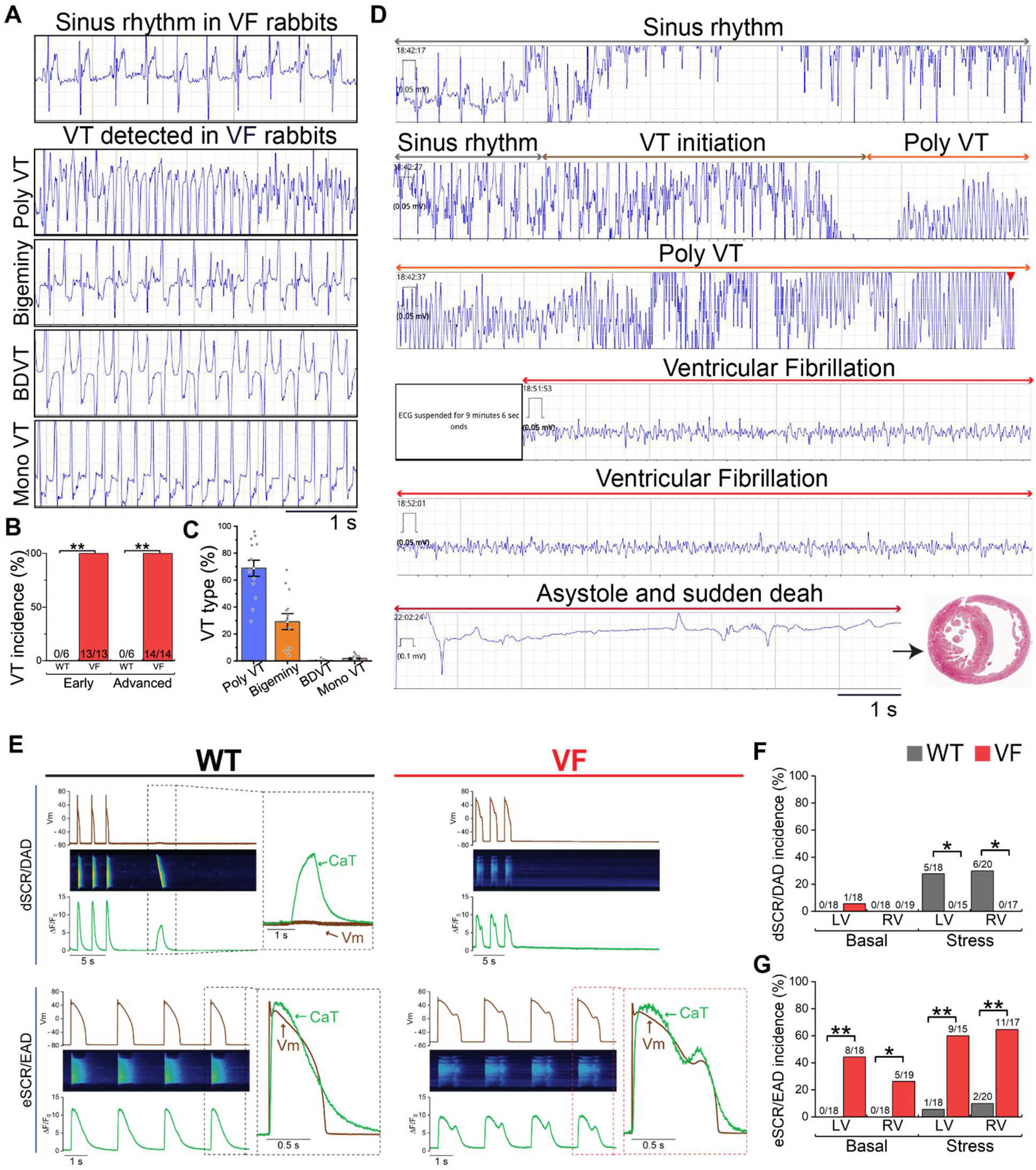
Lethal ventricular tachyarrhythmias and the underlying cellular mechanism. **A**. Representative traces of spontaneous ventricular arrhythmias in conscious VF rabbits: polymorphic ventricular tachycardia (poly VT), bigeminy, bidirectional ventricular tachycardia (BDVT), and monomorphic ventricular tachycardia (mono VT). **B**. VT incidence in WT and VF rabbits from early and advanced stage. N as indicated. **C**. Frequencies of different types of VTs recorded in each VF rabbit. N = 13 rabbits. **D**. Prolonged ventricular arrhythmia episode recorded prior to sudden death in a 14-week-old VF rabbit. MTS staining of a cross section of the heart is consistent with the early stage of disease progression. **E**. Representative simultaneous recordings of membrane potential (Vm) and confocal Ca^2+^ transients (CaT) in WT (left) and VF (right) cardiomyocytes under catecholamine stress during the early stage. Upper traces: Vm; middle: confocal CaT linescan image; lower: CaT fluorescence intensity profile. Insets show superimposed Vm and CaT signals from boxed regions. Upper panels illustrate diastolic spontaneous Ca^2+^ releases (dSCRs) and delayed afterdepolarizations (DADs); lower panels illustrate early spontaneous Ca^2+^ releases (eSCRs) and early afterdepolarizations (EADs). **F-G**. Incidence of dSCR/DADs (F) and eSCR/EADs (G) under basal conditions and catecholaminergic stress. Sample sizes of cardiomyocytes indicated in columns. For basal WT LV and RV, basal VF RV, stress WT LV and RV, stress VF RV, single cardiomyocytes were isolated from N = 6 rabbits each group; For basal VF LV, N = 8 rabbits; For stress WT RV, N = 7 rabbits. Chi-square or fisher exact test between stage and side matched WT and VF, *P < 0.05, **P < 0.01.

To elucidate the cellular basis of electrical instability, we performed simultaneous membrane voltage and Ca^2+^ transient (Vm/CaT) recordings in the early stage VF cardiomyocytes and age-matched WT controls. In RyR2 mutant mouse models, diastolic spontaneous Ca^2+^ release (dSCR) and delayed afterdepolarizations (DADs) are the main cellular drivers of arrhythmia.^20^ In contrast, VF cardiomyocytes from the early stage did not exhibit increased dSCR or DADs under basal conditions or during catecholaminergic stimulation (isoproterenol 50 nM). Instead, these cells predominantly exhibited early spontaneous Ca^2+^ release events (eSCR) during the Ca^2+^ transient decay and early afterdepolarizations (EADs) during the action potential plateau (**Figure 2E-G**). eSCR/EADs occurred in 44% of LV and 26% of RV VF cells but were absent in WT controls. Catecholaminergic stress further increased their incidence (**Figure 2G**). These findings identify eSCR/EADs as the dominant early arrhythmogenic mechanism in rabbits.

### Serial echocardiography reveals preserved early function followed by deteriorated RV contraction and impaired LV filling

To characterize the structural and functional progression from the early stage (11-12 weeks) to the advanced stage (20-22 weeks), we performed serial echocardiographic evaluations in VF rabbits and age-matched WT controls (categorized into the same stage based on age). Structurally, rabbit hearts undergo substantial physiological growth during the study period, as indicated by an increase in whole-heart area under both short axis view (SAX) and four chamber view (4CH) (**Figure 3A-D**). Compared to WT controls, VF rabbits developed RV enlargement during the early stage that progressed to biventricular dilation in the advanced disease (**Figure 3, Figure S7A-D**), consistent with the gross pathological findings.

**Figure 3.**
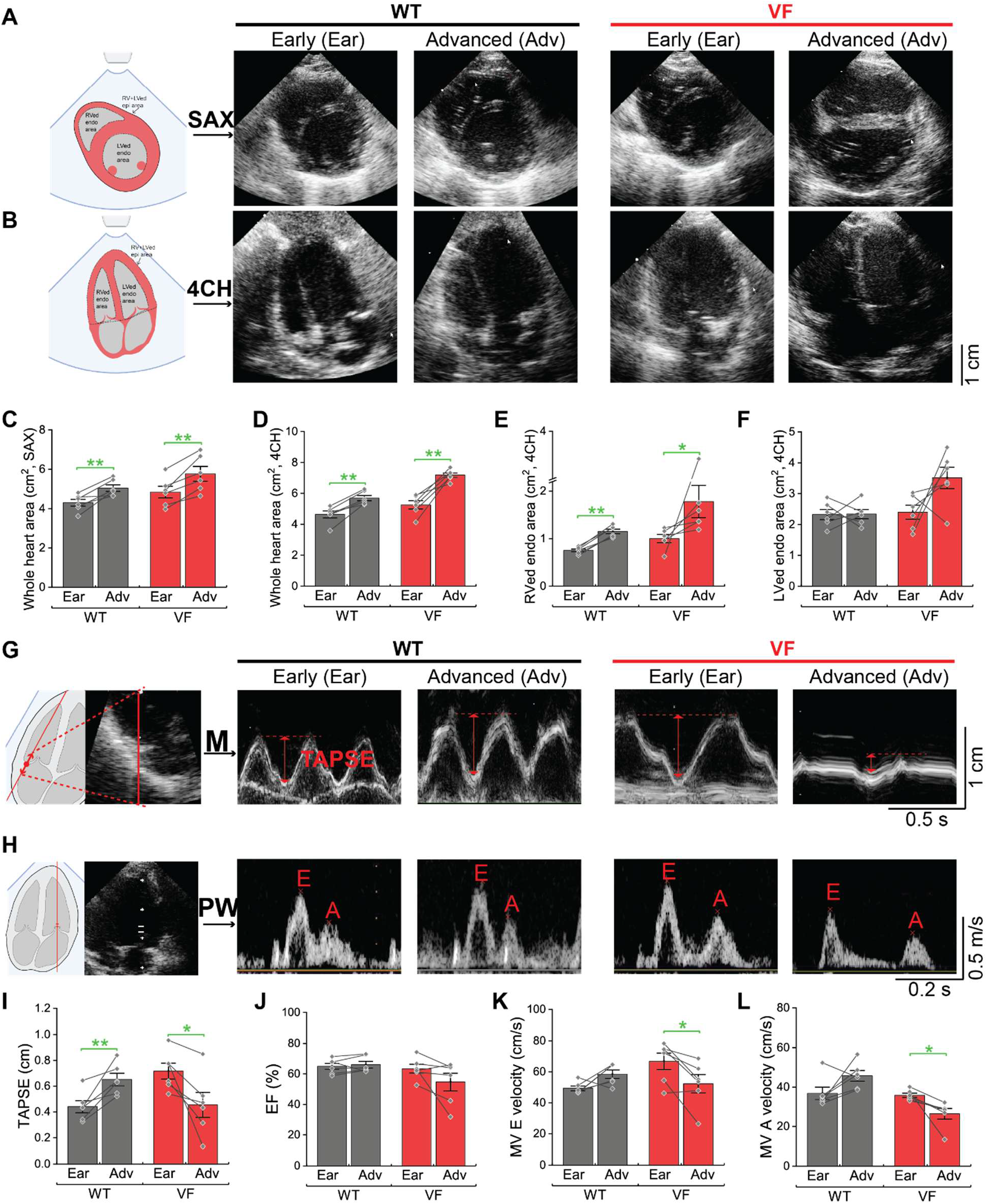
Progression of cardiac structure and function assessed by serial echocardiography. **A-F.** Cardiac structure. **A-B.** Representative echocardiographic images acquired in the short-axis (SAX, A) and four-chamber (4CH, B) views at the early (Ear) and advanced (Adv) stages. Measurements include whole heart area, end-diastolic right ventricular endocardial (RVed endo) area, and left ventricular endocardial (LVed endo) areas. **C-D**. Quantification of whole heart area in the SAX (C) and 4CH (D) views. **E**. RVed endo area in 4CH view. **F**. LVed endo area under 4CH view. **G-L**. Cardiac function. **G**. Representative M-mode echocardiographic images illustrating tricuspid annular plane systolic excursion (TAPSE) measurement for the RV. **H**. Representative pulsed-wave (PW) Doppler images showing mitral valve (MV) E-wave and A-wave measurements. **I**. Quantification of TAPSE for the RV. **J**. Quantification of LV ejection fraction (EF). **K-L**. Quantification of MV E velocity (K) and A velocity (L). N = 6 each group. Green asterisks indicate paired comparisons between Ear and Adv within genotype. *P < 0.05, **P < 0.01 by paired T test or signed rank test within genotype.

Despite early RV enlargement, RV systolic function was initially preserved, as measured by tricuspid annular plane systolic excursion (TAPSE, **Figure 3G, I, Figure S7E**) and RV systolic velocity (RV DTI s’, **Figure S7I-K**). Both early stage WT and VF animals showed similar RV DTI s’, indicating preserved RV contraction velocity (**Figure S7K**). TAPSE was modestly greater in the early stage VF hearts than in WT (**Figure 3, Figure S7E**), consistent with the larger RV chamber size. Importantly, as WT hearts grew, TAPSE increased in parallel with RV size growth. In contrast, VF rabbits exhibited a progressive decline in TAPSE despite continued RV dilation (**Figure 3G,I**), indicating worsening RV systolic dysfunction. This functional deterioration was further supported by reduced RV DTI s’ in advanced stage animals (**Figure S7K**).

LV systolic function remained largely preserved throughout disease progression, although 2/6 animals displayed ejection fraction below 45% (**Figure 3J, Figure S7F**), consistent with the heterogeneous clinical presentation of ACM.^21^ In contrast, LV diastolic function progressively deteriorated, as reflected by declining mitral E-wave and A-wave velocities (**Figure 3H,K-L**). These changes indicate worsening LV filling during disease progression and may reflect the functional consequences of chamber remodeling and compression from RV enlargement.

### RyR2 dysfunction triggers early adaptive excitation-contraction remodeling to preserve SR load but progresses to SR Ca^2+^ depletion

To determine the cellular mechanisms underlying progressive mechanical dysfunction, we next characterized excitation-contraction coupling (ECC) remodeling at both the cellular and molecular levels. At the cellular level, early stage VF cardiomyocytes isolated from both the LV and RV showed normal CaT amplitude (**Figure 4A,B**) and SR Ca^2+^ load (**Figure 4C**) compared to age-matched WT controls. Ca^2+^ release kinetics were slower in early stage VF myocytes, which showed prolonged time-to-peak (**Figure 4D**), reduced upstroke velocity (**Figure 4E**) and slower decay time (**Figure 4F**). At the molecular level, early stage VF hearts showed downregulation of all major Ca^2+^ handling proteins: RyR2, SR Ca^2+^ pump (SERCA2), Na^+^/Ca^2+^ exchanger (NCX1) and the α-subunit of the L-type Ca^2+^ channel (Ca_V_1.2) (**Figure 4G-K**). Lower RyR2 expression likely contributes to preventing SR Ca^2+^ depletion by limiting RyR2-mediated leak, as we have previously proposed.^22^ Downregulation of NCX1 may have a similar effect by preventing Ca^2+^ extrusion. Reduced Ca_V_1.2 and SERCA2 are likely to contribute to maintaining balanced Ca^2+^ fluxes across membranes. Overall, these adaptations may contribute to the slower intracellular Ca^2+^ transient dynamics in VF myocytes.

**Figure 4.**
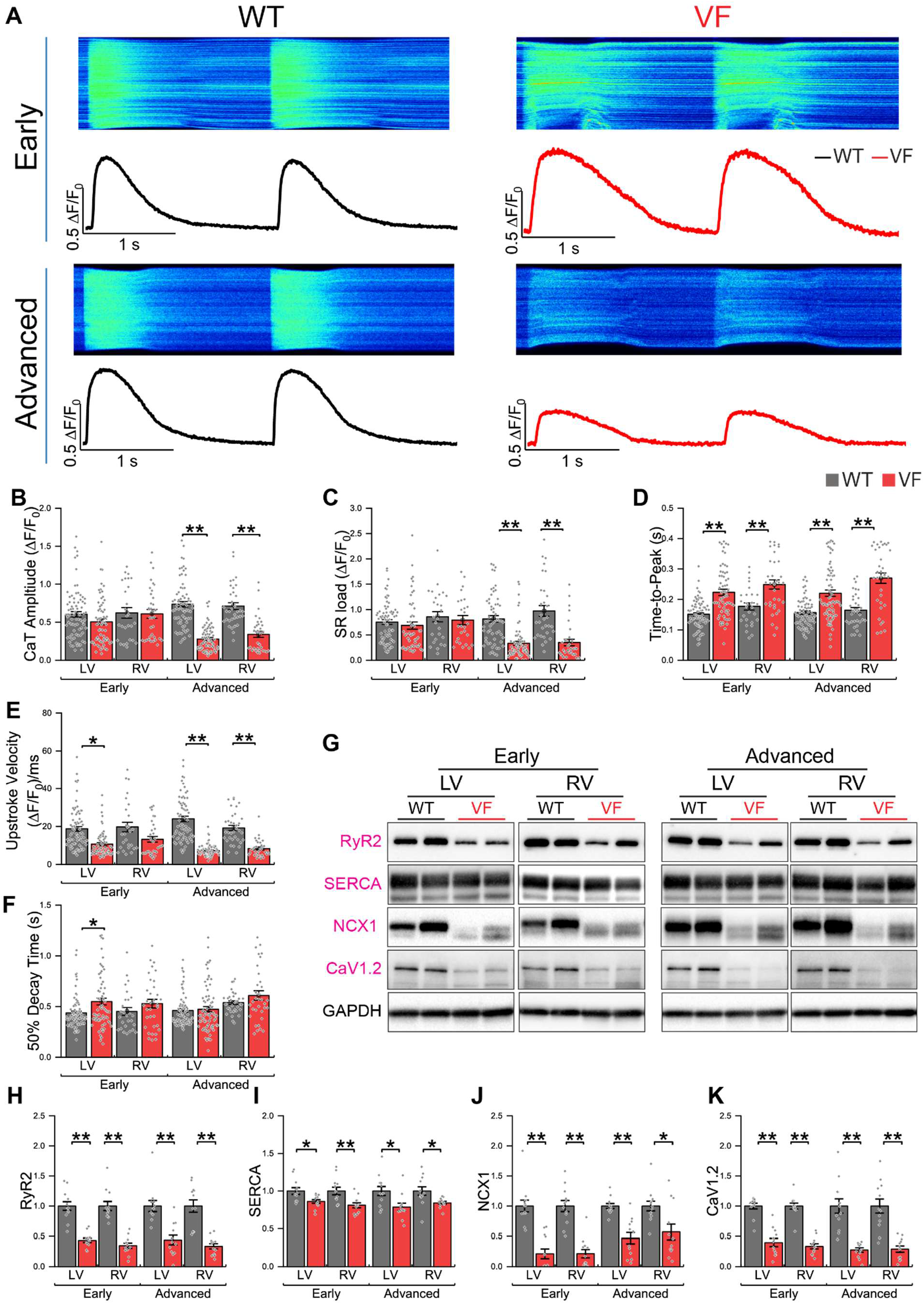
Early adaptive remodeling of excitation-contraction coupling in VF rabbit cardiomyocytes. **A**. Representative Ca^2+^ transient (CaT) recordings in isolated rabbit cardiomyocytes from WT (left) and VF myocytes (right) paced at 0.5 Hz. Upper: Early stage; lower: advanced stage. **B-F**. Quantification of CaT properties, including amplitude (B), SR Ca^2+^ load (C), time to peak (D), upstroke velocity (E), and 50% decay time (F). n for Early LV: WT = 74-77 cells, VF = 52-65 cells; n for Early RV: WT = 26-27 cells, VF = 26-34 cells. n for Advanced LV: WT = 59-70 cells, VF = 69-74 cells; n for Advanced RV: WT = 31-35 cells, VF = 33-34 cells. Single cardiomyocytes were isolated from 4 rabbits each group. *P < 0.05, **P < 0.01 by rabbit-level hierarchical analysis**. G-K**. Representative immunoblots and quantification of excitation-contraction coupling (ECC) proteins. SERCA2a, sarcoplasmic reticulum Ca^2+^ ATPase; Ca_V_1.2, L-type Ca^2+^ channel; NCX1, Na^+^/Ca^2+^ exchanger. N = 12 hearts per group. *P < 0.05, **P < 0.01; two-tailed t-test or Mann-Whitney test between stage and side matched WT and VF.

As disease progressed, advanced VF cardiomyocytes developed a failing phenotype with reduced Ca^2+^ transient amplitude and SR load (**Figure 4A-C**). The changes in Ca^2+^ release kinetics and ECC protein expression observed in the early stage were also present in the advanced stage (**Figure 4D-K**). Furthermore, both LV and RV showed similar ECC remodeling at the cellular and molecular levels (**Figure 4B-K**). Together, these findings indicate that chronic RyR2-mediated Ca^2+^ leak initially elicits adaptive remodeling that preserves SR Ca^2+^ load together with slower Ca^2+^ dynamics but ultimately progresses to SR Ca^2+^ depletion.

### Single-nucleus transcriptomics predicted cardiomyocyte-initiated multicellular remodeling in the early stage

To define the molecular pathways underlying the structural remodeling, we performed single-nucleus RNA sequencing (snRNA-seq) on LV and RV samples from early and advanced stage hearts (**Figure 5A**). After stringent quality control, ∼237,000 high-quality nuclei were retained and classified into nine major cell populations, including cardiomyocytes (CM), fibroblasts (FB), adipocytes (AD), and vascular, immune, and neuronal lineages (**Figure 5A, Figure S8, Figure S9**). The cell type distribution in WT hearts (40% CM, 26% FB, 0.28% AD) (**Figure 5A**) closely resembled that of the human myocardium,^23^ supporting the translational relevance of this model.

**Figure 5.**
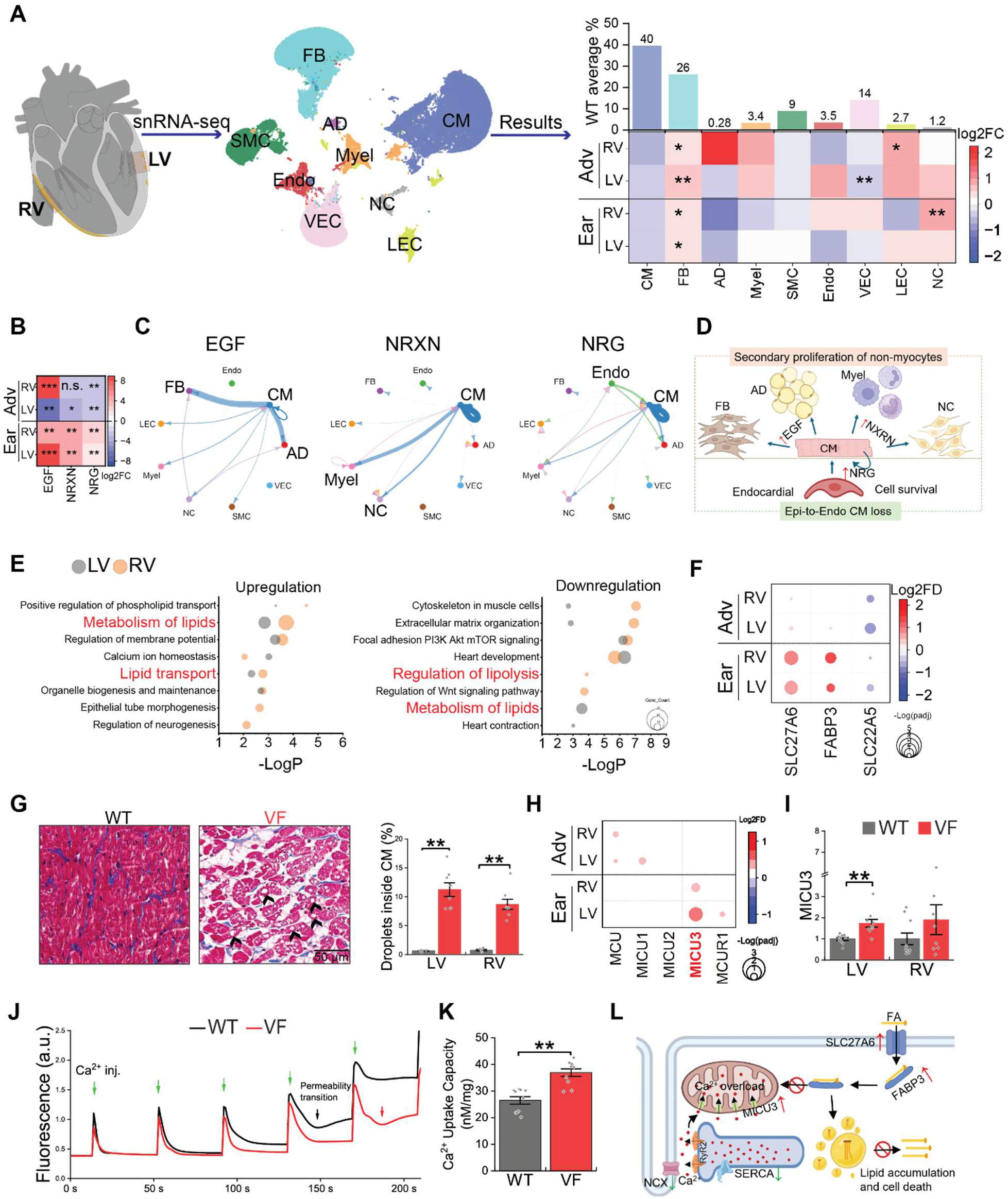
Single-nucleus RNA sequencing reveals early CM-centered multi-cellular remodeling and mitochondrial-associated lipid metabolic dysfunction. **A**. snRNA-seq experimental design and the identified cell types. Left: Nuclei isolated from LV and RV free wall were used for sequencing. Middle: UMAP embedding of 237,710 nuclei identified nine cell populations: cardiomyocytes (CM), fibroblasts (FB), adipocytes (AD), myeloid cells(Myel), smooth muscle cells (SMC), endocardial cells (Endo), vascular endothelial cells (VEC), lymphatic endothelial cells (LEC), and neuronal cells (NC). Right upper: relative cell-type proportions in WT hearts. Right lower: heatmap of Log2(fold change, FC) changes in VF hearts compared to WT controls. Ear: early stage; Adv: advanced stage. Red, increased; blue, decreased. N = 4 rabbits each group, except for N = 3 for the Adv VF rabbits. *P < 0.05, **P < 0.01; two-tailed t-test or Mann-Whitney test between stage and side matched WT and VF. **B**. Heatmap showing uniquely upregulated signaling pathways identified in CellChat database. Color scale represents log2FC in cell-cell communication strength. Dark red indicates pathways uniquely present in VF hearts. *Adjusted P < 0.05, **P < 0.01; ***P = 0 (present only in VF). **C**. Circle plots illustrating predicted significantly upregulated cell-cell communication pathways, including *EGF*, *NRXN*, *NRG* signaling in early stage VF LVs. Line thickness represents increased interaction strength (VF − WT). Edges color indicates source cell type. Arrows indicate signaling direction. **D**. Predicted cell-cell communication paradigm in RyR2-induced cardiomypathy. Early cardiomyocyte damage during the early triggers paracrine signaling that engages other cell populations as the disease progresses. **E**. Bubble plots of pathway enrichment for upregulated (left) and downregulated (right) differentially expressed genes (DEGs) identified in CM in the early stage. Bubble size indicates gene count. LV and RV pathways are color-coded by gray and orange as indicated. **F**. Expression changes of representative lipid metabolism genes. Color indicates log2FC; bubble size reflects -log(adjusted P value). **G**. Representative MTS images (left) and quantification (right) of intracellular lipid droplet accumulation in cardiomyocytes from the early stage (black arrows indicating big droplets). **H**. Bubble plot of DEGs associated with mitochondrial Ca^2+^ uptake. **I**. *MICU3* mRNA expression by qPCR. **J**. Representative traces of Ca^2+^ uptake by freshly isolated mitochondria from WT and VF LV in the early stage. Green arrows indicate sequential Ca^2+^ boluses (15 μmol/L each). VF mitochondria display enhanced cumulative uptake and delayed mitochondrial permeability transition pore (mPTP) opening, as indicated by black arrow in control and red arrow in VF group. **K**. Quantification of total mitochondrial Ca^2+^ uptake retention capacity prior to mPTP opening (N). **L**. Predicted mechanistic link between RyR2-mediated Ca^2+^ leak, mitochondrial dysfunction, and lipid accumulation. H-K n = 10 per group. *P < 0.05, **P < 0.01; two-tailed t-test or Mann-Whitney test.

In VF hearts, cell type analysis revealed a marked expansion of fibroblasts during the early stage, along with a nonsignificant trend toward reduced cardiomyocyte representation before overt extracellular fibrosis was detected (**Figure 5A**). In the advanced stage, fibroblast abundance did not significantly increase further (**Figure 5A**) despite extensive fibrosis (**Figure 1**), suggesting that fibroblast populations transition to active collagen deposition. Four fibroblast subpopulations were identified (FB1 -FB4, **Figure S10, Figure S11**), among which FB2 selectively expanded and acquired a collagen-producing phenotype. FB2 expansion occurred earlier and to a greater extent in the RV than in the LV (**Figure S10B,C**), paralleling the spatial progression of fibrofatty remodeling.

Adipocytes showed a tendency for population growth in the RV during the advanced disease (**Figure 5A**) but exhibited minimal transcriptional remodeling (**Figure S12**). This suggests that adipogenesis is likely secondary to cardiomyocyte remodeling rather than a primary disease driver. Additional increases in myeloid, lymphatic endothelial, and neuronal populations (**Figure 5A**) suggest coordinated inflammatory, vascular, and neurogenic responses.

Cell-cell communication analysis predicted injured cardiomyocytes as the dominant signaling hub during the early stage (**Figure 5B-D, Figure S13, Figure S14**). The predominant proliferative and remodeling pathways, including *EGF*^24^ and *NRXN*,^25^ were transiently activated during the early stage before overt structural remodeling, with increased paracrine signals from cardiomyocytes to fibroblast, adipocytes, neuronal and myeloid cells (**Figure 5B-C**). These pathways largely subsided once advanced remodeling became established, suggesting that the early stage represents the critical window during which stressed cardiomyocytes orchestrate multicellular remodeling. Among these pathways, neuregulin (*NRG*) signaling, a well-established cardioprotector,^26^ was selectively enhanced through both cardiomyocyte autocrine and endocardial-to-cardiomyocyte paracrine signaling, suggesting an adaptive response that may contribute to the characteristic epicardial-to-endocardial progression of disease (**Figure 5C-D**).

### Transcriptomic changes in cardiomyocytes suggest a link between SR Ca^2+^ leak and mitochondrial Ca^2+^ uptake

Although the remodeling observed in VF hearts resembled ACM, canonical signaling pathways involving desmosomal proteins and adipogenic markers were largely unchanged in cardiomyocyte transcriptomes (**Figure S15**). This was also verified by Western blots (WBs) and quantitative polymerase chain reaction (qPCR) on key molecules (**Figure S16**). Instead, pathway enrichment analysis identified lipid metabolism as the dominant transcriptional signature of early-stage cardiomyocytes, encompassing both upregulated and downregulated gene sets (**Figure 5E-F, Figure S17**). This metabolic reprogramming was characterized by increased expression of *SLC27A6*, a major cardiac fatty acid (FA) transporter, and *FABP3*, a cytosolic FA-binding protein, consistent with enhanced FA uptake and intracellular trafficking. This was coupled with downregulation of *SLC22A5*, which mediates mitochondrial FA import via carnitine transport, suggesting impaired FA utilization.^27^ Consistent with this transcriptional profile, lipid droplet-like vacuoles accumulated in early stage VF cardiomyocytes, occupying ∼10% of the cell compared with <1% in controls (**Figure 5G**). Together, these findings suggest early lipid metabolic reprogramming characterized by increased FA uptake coupled with impaired mitochondrial utilization, leading to intracellular lipid accumulation.

We next examined whether mitochondrial Ca^2+^ transport was adaptively remodeled. Among components of the mitochondrial Ca^2+^ uniporter (*MCU*) and its regulatory subunits (*MICU1-3* and *MCUR1*)^28^, only *MICU3* was significantly upregulated in the LV during the early stage, as identified by snRNA-seq and confirmed by qPCR (**Figure 5H-I**). Because *MICU3* enhances mitochondrial Ca^2+^ uptake,^29^ we directly assessed mitochondrial Ca^2+^ handling. Isolated mitochondria from early stage tissue displayed enhanced Ca^2+^ uptake and retention capacity, along with delayed opening of the mitochondrial permeability transition pore (mPTP) (**Figure 5J-K**). Mitochondria occupy ∼30% of cardiomyocyte volume;^19^ hence, this augmented Ca^2+^ buffering capacity could mitigate cytosolic Ca^2+^ overload. Consistent with this interpretation, canonical Ca^2+^-dependent stress signaling pathways, including CaMKII activation and calpain-mediated junctophilin-2 cleavage, remained largely inactive (**Figure S18, Figure S19**) despite persistent SR Ca^2+^ leak and limited Ca^2+^ extrusion. This implies that cardiomyocyte injury does not necessarily result from global cytosolic Ca^2+^ overload. Collectively, these findings suggest adaptation in mitochondrial Ca^2+^ handling in response to chronic RyR2 dysfunction (**Figure 5L**).

### Early RyR2 inhibition suppresses arrhythmias and improves cardiac function

The transcriptomic and functional remodeling in VF hearts manifest during the early stage, before overt remodeling is evident. Therefore, we next tested whether early long-term RyR2 inhibition with *ent*-verticilide, a selective allosteric RyR2 modulator,^30,31^ could prevent arrhythmias and slow functional deterioration in VF rabbits. Ten VF animals were randomized into two cohorts, receiving *ent*-verticilide or vehicle via a subcutaneous osmotic pump. Treatment started at 12 weeks of age (early stage) and continued for two months until the advanced stage. All animals underwent continuous ECG monitoring and biweekly echocardiographic assessment (**Figure 6A**). Baseline VT burden was similar between groups before treatment started (**Figure 6B**). No SD occurred among *ent*-verticilide-treated animals (0/5), compared with 2/5 in the vehicle group (**Figure 6C**) and 32% across the entire VF rabbit colony (37/115, all untreated, **Figure S4A**). Cumulative VT burden significantly diverged between the *ent*-verticilide and vehicle groups by day 4 and remained significantly lower throughout the treatment in the *ent*-verticilide-treated animals (**Figure 6D-E**). These results support the reported anti-arrhythmic properties of *ent-* verticilide and RyR2 inhibition.^32,33^

**Figure 6.**
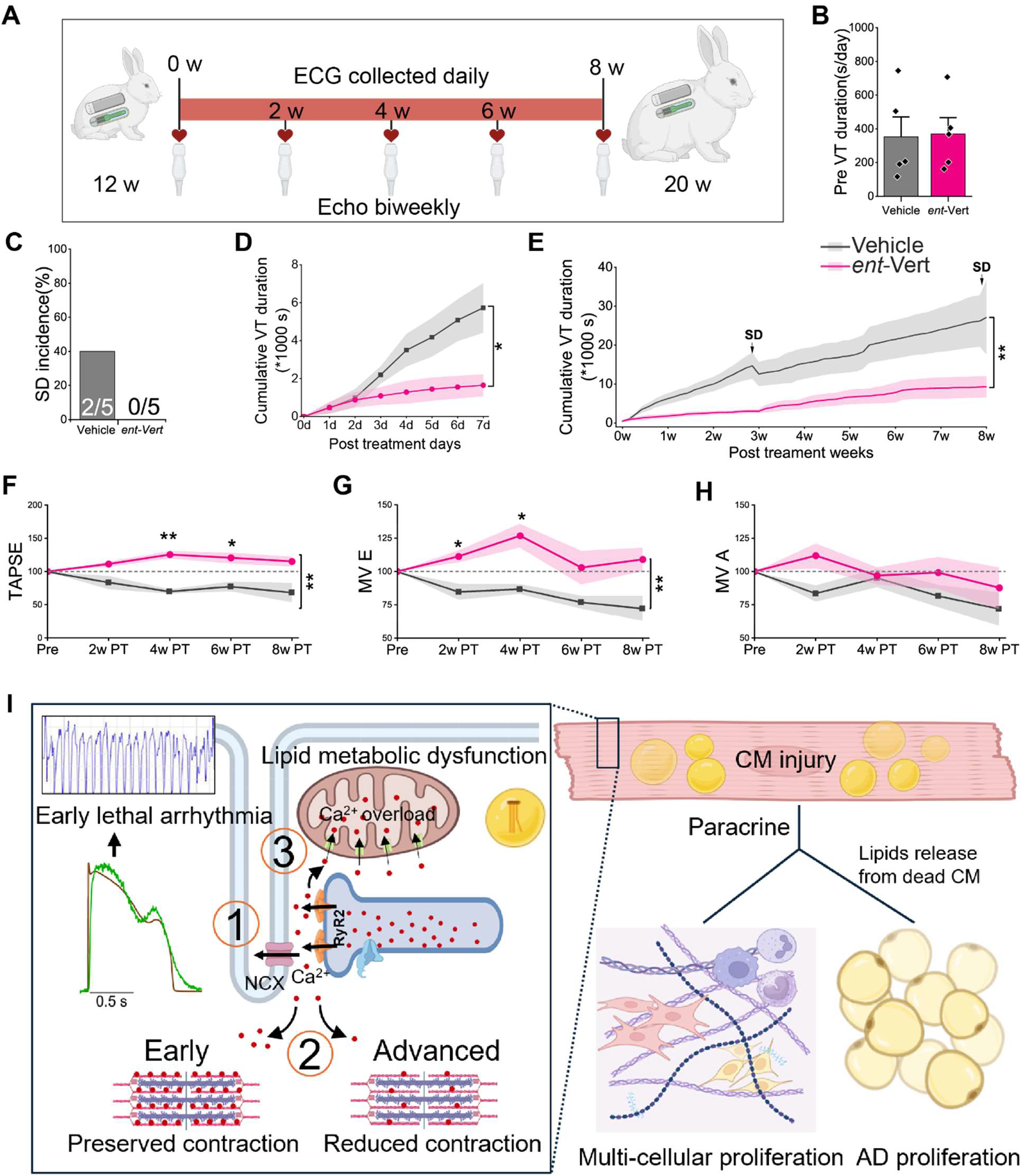
Therapeutic effects of selective RyR2 inhibitor *ent*-verticilide in ACM rabbit. **A**. Experimental design: Ten VF rabbits were implanted with telemetry ECG devices two weeks prior to treatment to establish baseline VT burden. At 12 weeks of age, animals were randomized to receive either *ent*-verticilide or vehicle via subcutaneous osmotic pumps for 8 weeks. Echocardiography was performed at baseline (prior to pump implantation) and biweekly to monitor disease progression. **B**. Average daily VT duration prior to treatment, showing no difference between the two groups. **C**. Incidence of SD during treatment: 2/5 in the vehicle group (similar to the 32% in untreated VF rabbits), 0/5 in the *ent*-verticilide group. **D-E**. Cumulative VT duration over the first 7-days(D) and entire 8-week treatment period (E). VT burden diverged by day 4 and remained significantly lower in the *ent*-verticilide group. Shaded area around the lines representsstandard error. SD: sudden death event in vehicle group. **F-H**. Echocardiographic changes relative to pretreatment: TAPSE (F), mitral valve E (G) and A (H) velocities. *ent*-Verticilide significantly improved diastolic and RV function: TAPSE increased at week 4 and 6 post-treatment (PT); MV E increased at weeks 2 and 4 post-treatment. For B–H: N = 5 rabbits per group. *P < 0.05, **P < 0.01 by Two way repeated ANOVA with mixed model (D-H) or t-test (B). Shaded area round the lines shows standard error. **I**. Summary of the proposed and predicted mechanism hypothesis. Chronic RyR2-mediated SR Ca^2+^ leak initiates a multiscale pathogenic cascade: (1) Increased SR Ca^2+^ leak promotes eSCRs, generating EADs and malignant ventricular arrhythmias. (2) During the early stage, adaptive remodeling of ECC machinery, including reduced RyR2 abundance and limited Ca^2+^ extrusion, helps preserve SR Ca^2+^ load and cardiac contraction. With disease progression, these compensatory mechanisms become insufficient, resulting in SR Ca^2+^ depletion and RV-dominant systolic dysfunction. (3) Persistent SR Ca^2+^ leak is accompanied by enhanced mitochondrial Ca^2+^ uptake, which may buffer cytosolic Ca^2+^ but is associated with impaired fatty acid metabolism and intracellular lipid accumulation, contributing to cardiomyocyte injury. Injured cardiomyocytes subsequently become a paracrine signaling hub, promoting expansion and remodeling of multiple cardiac cell populations. Lipids release following cardiomyocytes loss may cause adipocyte lipid sequestration and further contribute to adipocyte accumulation. Together, these processes link chronic RyR2 dysfunction to both electrical instability and ACM structural remodeling. *ent*-vert: *ent*-verticilide.

As expected, heart size and ventricular dimensions increased in both groups as rabbits grew during the monitoring period (**Figure S20A-H**). No significant differences in EF were observed between groups, although the study was not powered to exclude modest effects on LV systolic function (**Figure S20I**). Remarkably, *ent*-verticilide significantly improved RV systolic function and LV diastolic filling. TAPSE increased in rabbits receiving *ent*-verticilide (**Figure 6F**), showing a similar trend to WT rabbits as the heart grew (**Figure 3I**). In contrast, TAPSE declined in vehicle-treated rabbits. There was an overall significant increase in mitral valve E velocities, although not significant in MV A velocity, indicating improved LV filling and diastolic function (**Figure 6G-H**). Histological analysis showed the presence of fibrofatty infiltrations in all *ent*-verticilide treated rabbits, with similar extent as vehicle group (**Figure S21**). These data suggest that, while *ent*-verticilide does not fully reverse RyR2-induced structural remodeling, it reduces ventricular arrhythmia burden and preserves cardiac mechanical function.

## DISCUSSION

Abnormal SR Ca^2+^ release through RyR2 is widely recognized as a major trigger of ventricular arrhythmias. Because of the multifaceted role of Ca^2+^ in the regulation of signaling pathways linked to structural remodeling and hypertrophy, the possibility of establishing a link between RyR2 dysfunction and cardiomyopathic remodeling is tantalizing. Interestingly, a growing number of studies have identified RyR2 mutations in patients with different forms of cardiomyopathy.^7,13,14^ ACM, in particular, was the first structural disease linked to RyR2,^7,16^ but this association has been contested because of insufficient mechanistic support.^34^ Therefore, it has remained unresolved whether persistent RyR2 dysfunction can initiate myocardial degeneration.^34,35^ Confirmation of this potential link could prompt a reassessment of the role of RyR2 dysfunction in a wide range of cardiac disorders arising either from primary mutations in RyR2 itself or which involve dysregulation of the channel.

Using a genetically engineered rabbit model carrying the severe gain-of-function RyR2-V2475F mutation, we demonstrate that chronic RyR2-mediated SR Ca^2+^ mishandling is sufficient to induce the major pathological features of ACM, including malignant ventricular arrhythmias, progressive cardiomyocyte loss, right ventricular-predominant fibrofatty remodeling, and sudden cardiac death.^1^ More importantly, our study establishes a temporal framework linking early RyR2 dysfunction to progressive myocardial remodeling and demonstrates that pharmacological RyR2 stabilization initiated during the early stage of the disease suppresses arrhythmias and preserves cardiac function. This is the first demonstration that a primary insult in RyR2 can drive a cardiomyopathic phenotype consistent with human disease. Our findings identify chronic RyR2-mediated Ca^2+^ mishandling as a central mechanism that couples electrical instability with structural remodeling of the heart (**Figure 6I**). The timing and magnitude of the RyR2-mediated

dysfunction are clear distinctions with the traditional CPVT paradigm. Whereas stress triggers arrhythmogenic Ca^2+^ mishandling in CPVT, the presence of constitutively dysfunctional RyR2 channels (as in VF rabbits) may trigger a distinct cellular response that progresses to structural remodeling.

A major conceptual advance of this study is the definition of a disease trajectory initiated by SR Ca^2+^ mishandling. Previous RyR2-mutant mouse models largely reproduce stress-induced arrhythmias with little structural remodeling, leaving unresolved whether RyR2 dysfunction alone can drive cardiomyopathy. In contrast, the present rabbit model develops both malignant arrhythmias and progressive fibrofatty myocardial replacement that closely resembles human ACM. At the cellular level, RyR2 hyperactivity generated a distinct arrhythmogenic mechanism dominated by eSCR/EADs rather than the dSCR/DADs typically observed in rodents. This difference likely reflects fundamental species-specific Ca^2+^ handling. Whereas murine cardiomyocytes rely predominantly on SERCA-mediated SR Ca^2+^ recycling, rabbits and humans depend more heavily on sarcolemmal Ca^2+^ extrusion, shifting the vulnerable window for SCR from diastole to the decay phase of the Ca^2+^ transient.^19^ Chronic SR Ca^2+^ leak also triggered a coordinated adaptive remodeling of the ECC machinery that initially preserved SR Ca^2+^ content and mechanical function. As disease progressed, these compensatory mechanisms became insufficient, resulting in SR Ca^2+^ depletion, impaired Ca^2+^ transients, declining RV systolic function, and restricted LV filling. Together, these findings establish a direct mechanistic link between persistent RyR2 dysfunction and progressive cardiac dysfunction.

Our data provide mechanistic insight into how chronic RyR2 dysfunction drives myocardial remodeling. Rather than producing sustained cytosolic Ca^2+^ overload, cardiomyocytes likely adapted by simultaneously reducing the abundance of leaky RyR2 channels and enhancing mitochondrial Ca^2+^ uptake, thereby preventing global cytosolic Ca^2+^ overload despite persistent SR Ca^2+^ leak.^19,22,36–38^ This apparent adaptation was accompanied by metabolic remodeling, although whether enhanced mitochondrial Ca uptake directly contributes to altered fatty acid metabolism remains to be established. Consistent with cardiomyocytes being the primary site of injury, single-nucleus transcriptomics and cell-cell communication analyses suggested that stressed cardiomyocytes were the dominant signaling hub during the early stage. These observations support a model in which persistent RyR2 dysfunction initiates myocardial remodeling through primary cardiomyocyte injury, with fibrotic and adipogenic remodeling emerging as secondary responses to cardiomyocyte stress.

Our findings also have important implications for understanding ACM pathogenesis. Although the pathogenic role of RyR2 variants in ACM remains debated, increasing evidence suggests that abnormal RyR2 activity is a common downstream consequence of diverse genetic forms of ACM. Studies have shown that desmosomal deficiency enhances RyR2 activity through altered kinase signaling, producing increased SR Ca^2+^ leak and ventricular arrhythmias before structural remodeling becomes apparent.^2,4,39^ Our findings extend this concept by demonstrating that persistent RyR2 dysfunction alone is sufficient to drive progressive cardiomyopathy through desmosome-independent mechanisms. These results therefore support the broader concept that early Ca^2+^ mishandling represents a convergent disease mechanism contributing to myocardial degeneration across genetically distinct forms of ACM.

The translational implications of this study are twofold. First, the RyR2-V2475F rabbit provides a clinically relevant large-animal model that reproduces the defining features of human ACM, including RV-dominant remodeling, progressive epicardial-to-endocardial fibrofatty infiltration, preserved LV systolic function during early disease, and spontaneous sudden cardiac death. These characteristics likely reflect the close similarity of rabbit and human cardiac electrophysiology, excitation-contraction coupling, and myocardial architecture, underscoring the value of large-animal models for mechanistic studies and preclinical therapeutic evaluation.^1,40,41^ Second, continuous RyR2 inhibition initiated during the early stage markedly reduced ventricular tachycardia burden and improved RV systolic and LV diastolic function. Although established fibrofatty remodeling was not reversed over the treatment period, improvement of cardiac function despite persistent structural abnormalities suggests that RyR2 stabilization can modify disease progression even after remodeling has begun. These findings support RyR2 as a promising therapeutic target not only for suppressing ventricular arrhythmias but also for delaying functional deterioration in ACM.

Several limitations should be acknowledged. Although our data suggest enhanced mitochondrial Ca^2+^ uptake as an adaptive response linking chronic RyR2 dysfunction to metabolic remodeling, the molecular pathways connecting mitochondrial Ca^2+^ signaling to altered lipid metabolism require further investigation. In addition, cell-chat analysis predicted a CM-initiated signaling to other cell types that may contribute to subsequent multicellular remodeling. However, these have not been experimentally validated. Future studies using in vivo cell type-specific genetic or viral manipulation of candidate pathways and in vitro co-culture experiments will be important to determine the contribution of different cell types to the structural remodeling. The dosage of *ent*-verticilide in our study was limited by the osmotic pump capacity and solubility of the drug; however, even in these conditions, we confirmed the presence of *ent*-verticilide in rabbit plasma (not shown), and the drug markedly reduced ventricular arrhythmia burden and maintained cardiac function. Finally, RyR2-V2475F has not been directly linked to ACM in humans. Nonetheless, our study serves as a proof-of-concept supporting a direct link between RyR2 dysfunction and cardiomyopathy. This concept may be applicable to ACM-causing RyR2 variants as well as canonical forms of ACM in which RyR2 dysfunction is prominent during the early stage of the disease. Similarly, cardiomyopathic rabbits were homozygous for RyR2-V2475F. While homozygous RyR2 mutations are rare, there are cases documented in the literature.^42,43^ Our results should bring attention to the possibility of these patients showing symptoms of early cardiomyopathy. Additional studies will be necessary to establish a clinical link between homozygous RyR2 variants or specific severe gain-of-function variants and cardiomyopathy.

## Conclusion

Chronic RyR2-mediated SR Ca^2+^ leak is sufficient to initiate a multiscale pathogenic cascade that progresses from electrical instability and adaptive excitation-contraction coupling remodeling in cardiomyocytes to multicellular remodeling, and ultimately structural arrhythmogenic cardiomyopathy. By demonstrating that early pharmacological stabilization of RyR2 suppresses malignant arrhythmias while preserving cardiac function, this study establishes intracellular Ca^2+^ mishandling as both a fundamental driver of disease progression and a promising disease-modifying therapeutic target for arrhythmogenic cardiomyopathy.

## ACKNOWLEDGEMENTS

The authors thank the Center for Advanced Models for Translational Sciences and Therapeutics at the University of Michigan who contributed to the creation of the genetically engineered rabbit model, including Drs. Jifeng Zhang, Dongshan Yang and Jie Xu.

The authors also thank the Gene Expression Core at the Biotechnology Center (RRID: SCR_017757) and the Translational Research Initiatives in Pathology (TRIP) Laboratory at the University of Wisconsin-Madison for their services. The TRIP lab is supported by the UW Department of Pathology and Laboratory Medicine, UWCCC (P30CA014520) and the Office of The Director-NIH (S10OD023526).

The authors acknowledge Dr. Eric G. Schmuck of the Center for Biomedical Swine Research for providing assistance and technical support with telemetry and echocardiography assessments; Drs. Jim Stein and Farhan Raza of the Department of Medicine for their assistance with echocardiography interpretation; and Ms. Holly C. Dooge for technical support provided in support of in vivo experiments.

## SOURCES OF FUNDING

This work was supported by the National Institutes of Health (R01HL161070 to FJA; R01HL167195 to FJA and HHV; R01HL170144 to HHV) and start-up funds from the Department of Medicine (FJA). SE was supported by the National Institutes of Health-funded Graduate Training in Molecular and Cellular Pharmacology (T32GM141013). JZ was supported by the American Heart Association (26CDA1621322). AMG was supported by the University of Wisconsin-Madison Cardiovascular Research Center Summer Undergraduate Fellowship.

## DISCLOSURES

The authors report no competing interests.

## Supplementary Materials for

### Materials and Methods

#### General considerations

Animal use was approved by the University of Wisconsin-Madison School of Medicine and Public Health IACUC (M005944), which runs an animal program accredited by AAALAC. Animal husbandry was performed by specialized personnel following each institution’s standard procedures. All experiments performed at institutional cores were single blinded. Unless otherwise stated, all reagents were acquired from MilliporeSigma.

#### Generation of the RyR2-V2475F rabbit model

RyR2-V2475F (VF) rabbits were generated at the Center for Advanced Models for Translational Sciences and Therapeutics (CAMTraST), University of Michigan, using CRISPR/Cas9-mediated genome editing. To introduce the valine-to-phenylalanine substitution at the site equivalent to human RyR2 codon 2475, a guide RNA (sequence: 5′-GGGGTTTTGCCCAGATCACA-3′) was designed to target the Cas9 complex near the codon 2476 of the rabbit RyR2 gene. The donor vector contained the target point mutation along with synonymous silent mutations within the gRNA-recognition sequence to prevent recurrent cleavage following homologous recombination. The mixture of spCas9 mRNA, gRNA, and donor vector was microinjected into single-cell rabbit zygotes, which were subsequently transferred into pseudopregnant recipient females. Out of 23 kits born after a 30-day gestation period, four founder (F0) rabbits were confirmed as successful RyR2-V2475F knock-ins via PCR and Sanger sequencing. Founder rabbits (F0) were backcrossed with wild-type (WT) New Zealand White (NZW) rabbits (Charles River, strain code 052). F1 offspring were genotyped to confirm the presence of the targeted mutation. RyR2-V2475F rabbits were subsequently backcrossed with WT NZW rabbits for four additional generations prior to experimental use. WT littermates were used as controls. Age-matched WT and RyR2-V2475F rabbits of both sexes were used. Animals under 14 weeks were defined as the Early (Ear) stage, and those over 17 weeks as the advanced (Adv) stage.

#### Histology

For rabbits that died suddenly, hearts were harvested within 24 h postmortem. Hearts were rinsed in cold PBS to remove blood, and gross images (anterior and posterior views) were obtained prior to fixation. Whole hearts were fixed in 10% neutral buffered formalin for 48 h, washed three times (10 min each), and stored in 70% ethanol until processing. Transverse sections (∼2 mm thick) were obtained near the apex below the valves, embedded in paraffin, and stained with hematoxylin and eosin (H&E) and Masson’s trichrome staining (MTS) at the Translational Research Initiatives in Pathology (TRIP) lab and the Comparative Pathology Laboratory in UW-Madison. Histopathological diagnosis was performed by a veterinary pathologist in a blinded manner. Sections (5 μm) were used for H&E, MTS, and immunohistochemistry (IHC). Adipocytes were detected using anti-perilipin 1 (Invitrogen MA5-27861, 1:100). Slides were scanned at 40x using an Aperio AT2 digital slide scanner. Ventricular area and fibrofatty infiltration were quantified using Aperio ImageScope (Leica Biosystems), and fibrosis was expressed as a percentage of total tissue area. Intracardiomyocyte lipid droplets were quantified using ImageJ (Fiji).

#### Telemetry ECG

A Medtronic Reveal LINQ™ loop recorder was implanted subcutaneously in rabbits at the early stage. Anesthesia was induced with ketamine (20 mg/kg) and xylazine (5 mg/kg), and maintained with low-dose isoflurane (1-1.5%). The dorsal region between the scapulae was shaved, chemically depilated, and sterilized using povidone-iodine, 70% isopropanol, and chlorhexidine. A ∼1 cm incision was made, and a subcutaneous pocket (∼4–5 cm × 1 cm) was created for device placement. The incision was closed with sutures and tissue adhesive. Animals recovered within 20–30 min after anesthesia withdrawal. Continuous 24-hour ECG recordings were collected daily from conscious, freely moving rabbits. Arrhythmia events were manually annotated from collected ECG reports. Sustained arrhythmias were defined as lasting ≥5 s. Arrhythmia definitions: 1. Polymorphic VT (poly VT): continuously varying QRS morphology, axis, and amplitude; 2. Bigeminy: alternating sinus beats and premature ventricular complexes (1:1 pattern); 3. Bidirectional VT (BDVT): beat-to-beat alternation in QRS axis or morphology; 4. Monomorphic VT (mono VT): uniform QRS morphology and axis. Arrhythmia burden was quantified as the proportion of total arrhythmia duration attributable to each arrhythmia type.

#### Echocardiography

Transthoracic echocardiography was performed by registered cardiac sonographers at UW Health using a Philips iE33 Cart Diagnostic Ultrasound System equipped with an S12-4 transducer. Serial echocardiographic examinations were performed longitudinally in the same cohort of rabbits from the early to advanced disease stages. Rabbits were anesthetized with 1– 1.5% isoflurane and maintained on a heated platform at 37.0±0.5°C throughout imaging. Lead II ECG was obtained continuously via limb electrodes. The chest was shaved and residual hair was removed with a depilatory cream (Nair) before imaging. Cardiac chamber dimensions were measured in B-mode from short-axis and apical 4-chamber views. Left ventricular ejection fraction (LVEF) was determined from M-mode measurements obtained in the parasternal long-axis and short-axis views, with the values averaged across views. Tricuspid annular plane systolic excursion (TAPSE) was measured in the apical 4-chamber view using M-mode with the cursor aligned parallel to the longitudinal motion of the lateral tricuspid annulus. Right ventricular systolic tissue velocity (RV DTI s′) was measured in the apical 4-chamber view using Doppler tissue imaging, with the peak systolic myocardial velocity recorded at the lateral tricuspid annulus. Mitral inflow E- and A-wave velocities were measured using pulsed-wave Doppler in the apical 4-chamber view, with the sample volume positioned at the tips of the open mitral valve leaflets. All echocardiographic parameters were measured over at least 3 consecutive cardiac cycles.

#### Tissue homogenization

Left ventricular (LV) and right ventricular (RV) tissue homogenates were prepared as previously described ^44^. Briefly, hearts were pulverized in liquid nitrogen, suspended in homogenization buffer (0.9% NaCl, Tris-HCl 10 mM pH 6.8, 20 mM NaF, 2 µM leupeptin, 100 µM phenylmethylsulphonyl fluoride, 500 µM benzamidine, 100 nM aprotinin), homogenized using a Teflon pestle and centrifuged at 1,000 x*g* for 10 min at 4 °C. Supernatants were aliquoted and stored at -80 °C until used. Protein concentrations were determined using the Bradford method (Bio-Rad).

#### [^3^H]Ryanodine binding assays

Binding assays were performed as previously described ^22,45^. Binding reactions were prepared containing 100 μg of protein from heart homogenates. Ca^2+^ sensitivity curves were carried out in the presence of 0.2 M KCl, 20 mM Na-HEPES pH 7.4, 6.5 nM [^3^H]ryanodine (NET950, PerkinElmer), 1 mM EGTA and enough CaCl_2_ to set free [Ca^2+^] between 100 nM to 10 μM. The Ca^2+^/EGTA ratio for these solutions was determined using MaxChelator (WEBMAXCLITE v1.15; Temperature = 37 °C, pH = 7.4, Ionic = 0.2 N). Binding reactions were incubated for 2 h at 37 °C, then filtered through Whatman GF/B filters and washed three times with 5 mL of distilled water in a Brandel M24-R Harvester. Non-specific binding was determined in the presence of 20 μM unlabeled ryanodine (ALX-630-062, Enzo Life Sciences). [^3^H]ryanodine binding was measured by liquid scintillation using Bio-Safe II counting cocktail (RPI Research) in a TriCarb 4810TR scintillation counter. Hill’s equation was used to determine the maximum [^3^H]ryanodine binding and the EC_50_ in Origin 2025 (Origin Lab).

#### Western blotting

50 µg of tissue homogenate were suspended in Laemmlii buffer and separated by SDS-PAGE in 4-20% TGX precast gels (Bio-Rad). Proteins were then transferred to PVDF membranes using overnight wet transfer at 30 V. Membranes were probed with the following primary antibodies: anti-RyR [clone 34C] (1:2000; MA3-925, Thermo), pS2808-RyR (1:1000, custom-made),^46,47^ pS2030 (1:1000, custom-made),^22,47^ pS2814 (1:2000, A010-31, Badrilla), SERCA2 [clone 2A7-1] (1:1000; ab2861, Abcam), PLB [clone A1] (1:5000; A010-14, Badrilla), pS16-PLB (1:5000; A010-12, Badrilla), pT17-PLB (1:5000; A010-13, Badrilla), Cav1.2 (1:1000; ACC-022, Alomone), NCX1 [clone C2C12] (1:1000; MA3-926, Thermo), junctophilin-2 [clone H-3] (1:1000; sc-377086, Santa Cruz Biotechnology), PKP2 (1:1000, PA5-53144, Thermo), N-Cadherin [clone 3B9] (1:1000; 33-3900, Thermo), JUP (1:1000, Millipore Sigma, P8087), CaMKII (1:1000, ab181052, Abcam), pT286-CaMKII (phospho T286) (1:1000, ab32678, Abcam, Boston, MA), and GAPDH [clone 6C5] (1:10000; MAB374, Millipore). Secondary antibodies, used as appropriate, were goat anti-rabbit-HRP (1:1000 or 1:5000; 31437, Thermo) or goat anti-rabbit-HRP (1:1000 or 1:5000; 31463, Thermo). Membranes were developed using SuperSignal ECL reagent (Thermo) and imaged with a ChemiDoc MP apparatus (Bio-Rad). Band intensity was quantified with the ImageLab software (Bio-Rad).

#### RNA extraction and qPCR

RNA was extracted using the PureLink™ RNA Mini Kit (Thermo Fisher). cDNA synthesis was performed using iScript™ Reverse Transcription Supermix (Bio-Rad). Quantitative PCR was conducted using iTaq Universal SYBR Green Supermix on a CFX Duet system (Bio-Rad).

Primer sequences: PPARG: F: 5′-ACTAGGGACAGATGCGATGAG-3′; R: 5′-CATCCACTGCTCTCAACCAC-3′; MICU3: F: 5′-TTGCTAAAACATGGAAGTCAC-3′; R: 5′-TCTGTATAAGAAATCACACCTCT-3′.

#### Isolation of cardiac myocytes

Ventricular myocytes were isolated as previously described ^22^. Rabbits were sedated with 30 mg/kg ketamine and 5 mg/kg xylazine, heparinized (1000 U/kg), and fully anesthetized with 10 mg/kg propofol (Abbott). Hearts were mounted on a Langendorff and perfused for 5–10 min (30-40 mL/min, 37 °C) with normal Tyrode’s (NT) solution containing (mM) 138 NaCl, 4.4 KCl, 1 MgCl2, 24 HEPES, 11 glucose, 1.8 CaCl2, pH 7.4, aerated with 100% O2. Once blood was completely removed, perfusion was switched to nominally-free Ca^2+^ Tyrode’s solution for 5–10 min until the heart stopped contracting. The perfusion was then changed to 36 µM Ca^2+^ Tyrode’s solution supplemented with 200 U/mL collagenase type 2 (Worthington), 1 mg/mL BSA and, for 15-30 min until the heart became flaccid. The LV and RV were then separated and minced in collagenase solution. Myocytes were dispersed by gentle pipetting. The supernatant was collected and the remaining tissue was further digested in fresh collagenase solution. This process was repeated, every 3-5 min, six times. After gently dispersion, the cell-containing supernatants were centrifuged (100 x g, 30 s) and the myocytes were resuspended in Tyrode’s solution containing 1 mg/mL BSA and 36 µM Ca^2+^. The solution was replaced every 10–15 min to raise [Ca^2+^] to 1.8 mM in five steps. Cells were maintained in NT solution until used.

#### Patch clamp

Whole-cell patch-clamp experiments were conducted using either an Axopatch 200B amplifier with a Digidata 1322A digitizer or a MultiClamp 700B with a Digidata 1550B digitizer (Axon Instruments) at room temperature. Ca^2+^ transients were imaged during simultaneous action potential/Ca^2+^ transient (AP/CaT) recordings using an LSM800 confocal microscope with a sampling rate of 2.47 ms/line.

For Vm/CaT recordings, the bath solution contained (in mM): 140 NaCl, 4 KCl, 1.8 CaCl_2_, 1 MgCl_2_, 10 HEPES, 1.2 NaH_2_PO_4_, and 10 glucose, pH adjusted to 7.4 with NaOH. The pipette internal solution consisted of (in mM): 120 K aspartate, 8 KCl, 7 NaCl, 1 MgCl_2_, 5 MgATP, 0.3 Na_2_GTP, 10 HEPES, 10 glucose and 0.3 Fluo-4-pentapotassium salt (Invitrogen F14200), pH adjusted to 7.2 with KOH. Cells were current-clamped, and action potentials were elicited using an A-M Systems^TM^ 8-channel Stimulator (model 3800) with a pulse width of 2 ms. Cells were paced at 0.5 Hz for 40s followed by a subsequent 20 s resting periods to observe the diastolic spontaneous Ca^2+^ release. Under stressed condition with catecholamine, 50 nM isoproterenol was supplemented to the bath NT solution.

#### Ca^2+^ transients

Ca^2+^ transients elicited by field stimulation (2 ms pulses, 60-80V) and SR Ca^2+^ load were recorded as described in the literature.^47^ Myocytes were incubated for 5 min at 37 °C in 10 µM fluo-4 AM with 0.4% Pluronic F-127 (ThermoFisher) and then plated on laminin-coated glass-bottom dishes (MatTek) with NT solution. Fluorescence was excited at 488 nm and recorded at >505 nm. Longitudinal line-scans were recorded at 0.5 Hz field stimulation using an LSM800 confocal microscope (Carl Zeiss) with a 40x/1.2 water immersion objective at 2.47 ms/line. 10 mM caffeine was applied to measure the SR Ca^2+^ load. At least five Ca^2+^ transients were averaged from each condition. Ca^2+^ transients were analyzed using custom MATLAB 2025 (Mathworks) scripts.

#### Mitochondrial isolation and Ca^2+^ uptake measurement

Mitochondrial Ca^2+^ uptake was measured in mitochondria isolated from snap-frozen cardiac tissue, following a previously published protocol with minor modifications.^48^ Briefly, 50–70 mg of heart tissue was minced into small pieces on ice and digested in 10 mL of ice-cold mitochondrial isolation medium (MIM; 300 mM sucrose, 10 mM Na-HEPES, 0.2 mM EDTA, and 1 mM EGTA, pH 7.2) supplemented with trypsin (125 μg/mL) for exactly 2 min. Digestion was terminated by adding 10 mL of stopping buffer (MIM containing 1 mg/mL BSA and 650 μg/mL trypsin inhibitor). The suspension was gently inverted several times and allowed to settle for 3–5 min. The supernatant was discarded, and the tissue pellet was resuspended in 4 mL MIM supplemented with BSA. Samples were homogenized (∼11 strokes) until no visible tissue fragments remained. The homogenate was transferred to a 15 mL tube and centrifuged at 600 × g for 10 min at 4°C to pellet nuclei and unbroken cells. The supernatant was then transferred to a new tube and centrifuged at 8,000 × g for 15 min at 4°C to pellet mitochondria. The mitochondrial pellet was retained and washed twice with 10 mL ice-cold MIM supplemented with BSA. For each wash, the pellet was resuspended and centrifuged at 8,000 × g for 15 min at 4°C. After the final wash, the mitochondrial pellet was resuspended in 100 μL ice-cold MIM and kept on ice. Protein concentration was determined using the Bradford assay. Mitochondria were used for experiments within 1 h of isolation.

For Ca^2+^ uptake measurements, 100 μg mitochondrial protein was added to 200 μL assay buffer containing 125 mM KCl, 20 mM K_2_HPO_4_, 20 mM HEPES, 1 mM MgCl_2_, 5 mM pyruvate, 5 mM succinate, and 5 mM malate (pH 7.2). Samples were gently mixed and incubated for 2 min at room temperature to allow mitochondrial energization. Calcium Green-5N (1 μL of 1 mM stock; Thermo Fisher, C3737) was added, and fluorescence was monitored using a CLARIOstar Plus plate reader (BMG Labtech) with excitation at 483 nm and emission at 530 nm. Fluorescence was recorded kinetically at 1 s intervals. Sequential Ca^2+^ pulses were delivered using automated injectors (5 μL of 0.6 mM CaCl_2_) at 20, 65, 110, 155, 195, 235, 275, and 315 s. The plate was shaken for 2 s at 200 rpm prior to each injection to ensure mixing.

Partial Ca^2+^ uptake for each pulse was quantified using a calibration curve to determine the fraction of Ca^2+^ not sequestered by mitochondria. Total Ca^2+^ uptake capacity was defined as the cumulative Ca^2+^ uptake across successive injections prior to permeability transition.

#### Single Nuclei Isolation and Processing

Nuclei were isolated from frozen transmural LV and RV free walls. Four biological replicates were used per group, except for VF hearts from the advanced stage (n=3) due to the high mortality.

Nuclei were isolated following a previously described protocol with modifications to accommodate this study ^49^. Briefly, 20–30 mg of frozen heart tissue was minced into 0.5–1 mm pieces on ice using a sterile scalpel blade. The tissue fragments were transferred to a 2 mL flat-bottom centrifuge tube containing a micro-stir rod and 1 mL of ice-cold lysis buffer (Nuclei EZ Lysis Buffer, Sigma, N3408) supplemented with 200 U/mL RNase inhibitor (RiboLock RNase Inhibitor, ThermoFisher, EO0382) and 50 U/mL DNase I (ThermoFisher, EN0523). Lysis was carried out by placing the tube on a magnetic stir plate at 100 RPM for 5 minutes on ice. Following the first lysis, the supernatant was transferred to a 15 mL Protein LoBind tube (Eppendorf, 30122216) containing 6 mL of ice-cold wash buffer (1× PBS, pH 7.4, 2% bovine serum albumin, 200 U/mL RNase inhibitor, and 50 U/mL DNase I). To maximize nuclei yield, the lysis and wash steps were repeated three times. The pooled lysate was sequentially filtered through 70 µm, 40 µm, and 20 µm cell strainers (pluriStrainer®, 43-50070, 43-50040, and 43-50020, respectively) to remove debris. The filtered nuclei suspension was transferred to a fresh 15 mL LoBind tube and centrifuged at 600×g for 5 minutes at 4°C. The pellet was resuspended in 200 μL of ice-cold wash buffer and gently pipetted 7–10 times to minimize nuclear aggregation.

Nuclei were stained with DAPI (Thermo Fisher Scientific, D3571) and Trypan blue (Thermo Fisher Scientific, 15250061) to assess the nuclei quality and quantity. Quantification was performed manually with a hemocytometer and automatically using Bio-rad TC20 Automated Cell Counter. Immediately after isolation, nuclei were fixed using the Evercode™ Nuclei Fixation v3 kit (Parse Bioscience, ECFN3300) according to the manufacturer’s instructions and stored at −80°C. Frozen nuclei samples were sent to the University of Wisconsin Biotechnology Center (UWBC) for barcoding and library preparation using the Evercode™ WT v3 kit (Parse Bioscience, ECWT3300). The barcoding process was performed using Parse Bioscience’s split-pool combinatorial indexing method. Prior to barcoding, thawed nuclei were visually inspected under a microscope to assess integrity and counted using the Countess II automated counter (Life Technologies). Quality control of the final cDNA libraries was performed using the Bioanalyzer High Sensitivity DNA Analysis kit (Agilent). Libraries were sequenced on an Illumina HiSeq platform with a target sequencing depth of 20,000 reads per nucleus.

#### Data Processing, Quality Control, and Clustering

FASTQ files were uploaded to Trailmaker (Parse Biosciences, https://app.trailmaker.parsebiosciences.com). Each sample was mapped to the rabbit reference genome OryCun2.0 using Trailmaker. After barcode correction, read alignment, and deduplication, quantified transcripts were used to generate a cell-by-gene count matrix for downstream analysis. The sequencing saturation was 24.7%, with base quality scores (Q30) for each barcode (BC) as follows: BC1 (96.0%), BC2 (96.2%), BC3 (97.2%), cDNA (96.5%), and polyN (96.1%). The proportion of reads and transcripts originating from rabbit nuclei were 74.5% and 74.3%, respectively.

To remove empty droplets and debris, EmptyDrops (implemented in Trailmaker) was applied with a false discovery rate (FDR) threshold of 0.01. Nuclei were filtered based on the number of detected genes and transcripts, using default spine fit-defined thresholds. Cells with mitochondrial gene content >1% were excluded. Doublet removal was performed using the scDblFinder with a probability of being a doublet < 0.4. Additionally, single nuclei with < 500 detected transcripts were removed from the dataset.

For batch effect correction and integration, the Harmony method in Seurat was applied using 2,000 highly variable genes (HVGs) and log-normalization. Principal component analysis (PCA) was conducted, and an elbow plot indicated a sharp decline in variance explained after the first 9 PCs, which together accounted for 73% of the total variance. Therefore, 9 PCs were selected for dimensionality reduction. Uniform Manifold Approximation and Projection (UMAP) was used for embedding visualization with a minimum distance of 0.1 and the Euclidean distance metric. Louvain clustering was performed at a resolution of 0.1. Clusters with highly similar differentially expressed genes were merged, based on shared marker genes and expression profiles.

Nuclei were classified into cell types using *ScType-Heart-Human*. For clusters that remained unassigned, manual annotation was performed based on the top 5 marker genes, with gene enrichment analysis conducted using *CellMarker 2024* via *Enrichr* (https://maayanlab.cloud/Enrichr/enrich). To identify cell-state subpopulations, fibroblasts (FBs) were further subclustered. For the FB subset, we selected 9 PCs and applied Louvain clustering with a resolution of 0.1.

#### Differential Abundance Analysis

We have 4 pairs matched groups, namely Early LV (Ear LV) from VF and the matched WT control, Ear RV from VF and WT, Advanced LV (Adv LV) from VF and WT, Adv RV from VF and WT. For each group, the abundance of each cell type was calculated as the ratio of the number of cells of that type to the total number of identified cells within the group. Only stage- and ventricle-matched cell types were included in the abundance comparison. Changes in cell type abundance were assessed by calculating the fold change (FC) between VF and matched WT controls. A fold change greater than 1 indicates an increased abundance in the VF group. To improve visualization, FC values were log2-transformed, where positive Log2FC values indicate higher abundance in the VF hearts, and negative values indicate lower abundance compared to matched WT controls. Statistical significance was assessed using the Chi-square test or Fisher’s exact test in SigmaPlot 14.0.

#### Differential Gene Expression and Variance Analysis

Differential expression analysis between different cell clusters was performed using the presto implementation of the Wilcoxon rank sum test and area under the receiver operating characteristic curve (auROC) analysis in Trailmaker. This was used to identify marker genes for each cell type or subcluster.

To compare gene expression between groups within a specific cell type, a pseudobulk limma-voom workflow (https://bioconductor.org/books/3.14/OSCA.workflows/segerstolpe-human-pancreas-smart-seq2.html#segerstolpe-comparison) was used. Briefly, pseudo-bulk differential expression sums the counts for all cells within a cluster for each sample and then uses standard differential expression methods designed for bulk RNA-seq. Differentially expressed genes (DEGs) were identified using the Wilcoxon rank sum test, with statistical significance defined as adjusted p-value (Padj) < 0.05 and |Log2FC| > 0.5. In this study, comparisons were exclusively conducted between stage- and ventricle-matched VF and WT samples.

#### Disease and Pathway Enrichment Analysis

Disease and pathway enrichment analysis was performed using Metascape^50^ (http://metascape.org) with the DEGs lists as input. Pathway enrichment was conducted separately for upregulated and downregulated genes within each cell type. Disease enrichment analysis was performed exclusively on DEGs identified in CMs using DisGeNet via Metascape. Both upregulated and downregulated DEG lists were used as input for disease association analysis.

#### Cell-cell communication analysis

Cell-cell interactions between assigned cell types in LVs and RVs from early and advanced stages were inferred using CellChat V2 in Rstudio according to the published tutorial protocol by Jin et al.^51^ The package and cellchat database is accessible at https://github.com/jinworks/CellChat.

Because CellChat relies on a human ligand–receptor interaction database, rabbit gene expression matrices were converted to human gene symbols using a rabbit-to-human ortholog mapping table from Ensembl. Only genes with valid human orthologs (59.61%,17636/29587) were retained. For genes mapping to the same human symbol, expression values were aggregated by summation. The resulting humanized count matrices were used to construct new Seurat objects for each condition.

Datasets were subset into VF and matched control WT groups, and CellChat objects were created using raw count matrices and cell-type annotations as grouping variables. The built-in human ligand–receptor database (*CellChatDB.human*) was used for downstream analysis.

Preprocessing included subsetting signaling genes (*subsetData*), followed by identification of overexpressed genes (i*dentifyOverExpressedGenes*) and ligand–receptor interactions (*identifyOverExpressedInteractions*). Communication probabilities between cell types were computed using the truncated mean approach (*computeCommunProb*, type = “truncatedMean”, trim = 0.05) based on raw expression data. Interactions from cell populations containing fewer than 10 cells were filtered out.

Cell–cell communication was further analyzed at the signaling pathway level using *computeCommunProbPathway*, and global interaction networks were summarized using *aggregateNet*. Individual CellChat objects were merged to enable direct comparison between VF and matched WT conditions.

Differences in total interaction number and interaction strength were quantified using *compareInteractions* and visualized using circle plots and heatmaps. Differential signaling pathways were identified using *rankNet*, with statistical testing enabled (do.stat = TRUE). For each pathway, contribution scores were extracted, and relative information flow was calculated as the proportion of total signaling contribution within each condition. Additional metrics, including fold change, dominance index, and pathway-specific interaction strength differences, were computed to characterize condition-specific signaling alterations.

For pathway-specific visualization: Edge thickness represents interaction strength of VF–WT differences; Edge color indicates source cell type; Arrows indicate signaling direction.

#### Study with ent-verticilide

*ent*-Verticilide was synthesized as previously described.^52–54^ *ent*-Verticilide was dissolved in a vehicle containing 50% dimethyl sulfoxide (DMSO) and 50% polyethylene glycol 400 (PEG 400) [https://alzet.com/formulating-the-solution/] and loaded into 2 mL osmotic pumps (ALZET, model 2ML4). Two osmotic pumps, each loaded with 25 mg *ent*-verticilide (50 mg in total), were implanted subcutaneously in the dorsal region of each rabbit to achieve the target dose. Rabbits also received a Medtronic Reveal LINQ™ loop recorder, as described earlier, for ECG monitoring. Four weeks after implantation, the pumps were surgically replaced with new pumps to maintain continuous drug delivery for the duration of the 8-week treatment period. Retrieved pumps were inspected to confirm complete or near-complete solution delivery. Based on the rabbit body weight, osmotic pump delivery rate and capacity, and *ent*-verticilide solubility, estimated dose range was of 0.6-0.7 mg/kg/day. Blood samples were collected during echocardiographic assessments to measure plasma concentrations of *ent*-verticilide as previously described.^31,32^

#### Statistical analysis

Numerical results and bar graphs are given as mean ± SEM with individual data points overlayed, unless otherwise noted. Statistical significance was determined at *p* < 0.05 using the tests indicated in each figure. In general, comparisons in animal level measurements between genotypes were done independently between matched WT and VF or Vehicle and *ent*-verticilide group, using t-test (parametric) or Mann-Whitney Rank Sum test (non-parametric); comparisons between proportions were done using fisher exact or chi-square test. Measurements in single cardiomyocyte were tested by rabbit level hierarchical analysis.^55^ All analyses, including additional tests for normal distribution and equal variance, as required, were carried out in SigmaPlot 14 (Systat Software) and R 4.5.2. Data management was done in Excel 365 (Microsoft) and Origin 2025 (Origin Labs). All test results can be found in supplementary table 1.

Additional information and reagents can be made available from the corresponding author upon reasonable request.

## Supplementary figures

**Figure S1.**
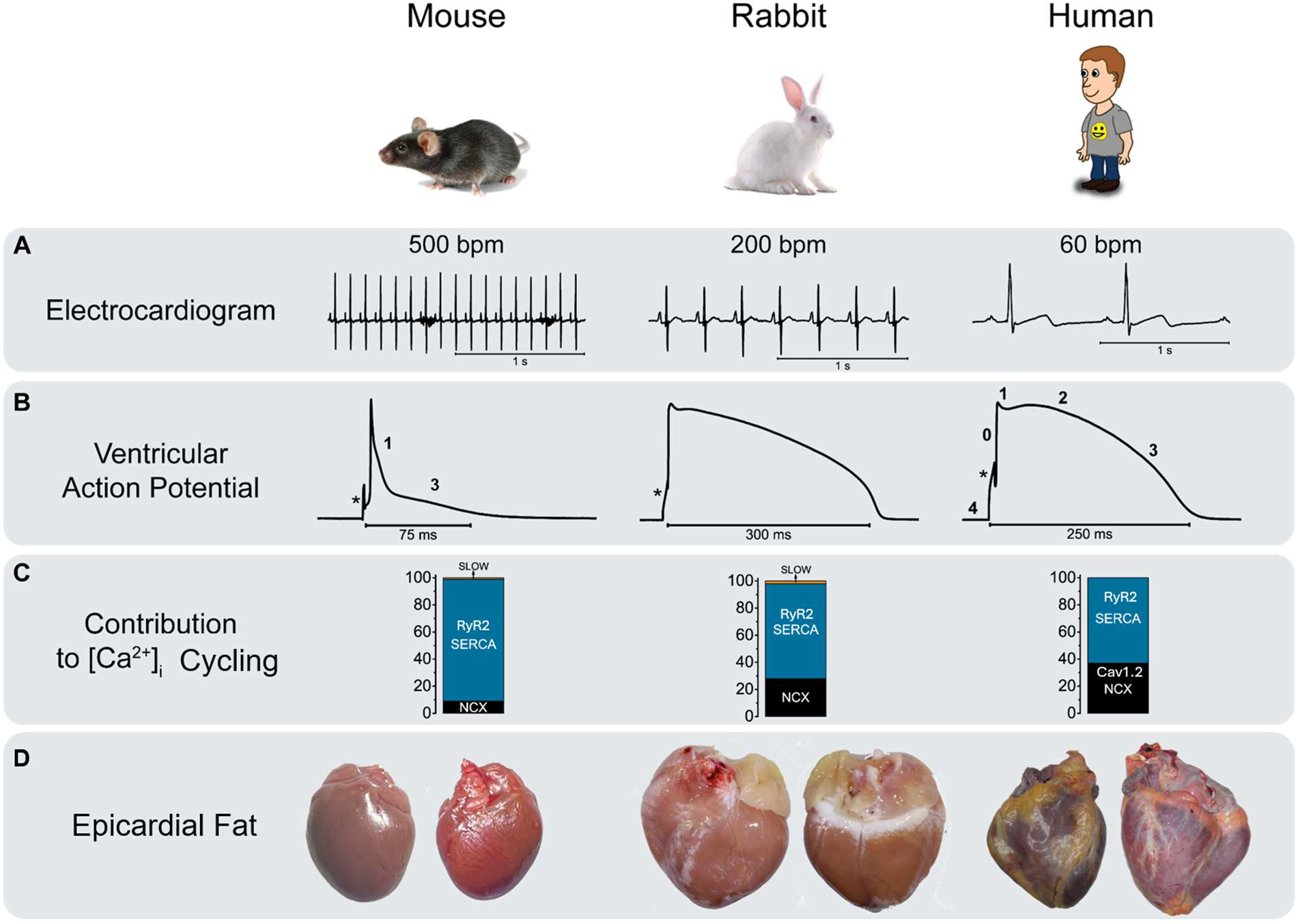
Species difference in electrophysiology, Ca^2+^ handling and myocardial architecture. **A-D.** Comparisons of surface ECG (A), ventricular action potential morphology (B), relative contributions to Ca^2+^ cycling (C), and normal epicardial fat distribution differences among mouse, rabbit and human (D). Rabbits exhibit an intermediate heart rate (∼200 bpm), between that of mice (∼500 bpm) and humans (∼60 bpm). Isolated rabbit ventricular cardiomyocytes display action potentials with a distinct plateau phase, resembling human action potentials and contrasting with the brief, triangular action potentials seen in mice (asterisks indicate stimulus artifact). During excitation–contraction coupling, cytosolic Ca^2+^ influx and efflux in human and rabbit myocytes is supported by approximately 30% sarcolemmal (LTCC and NCX) and 70% sarcoplasmic reticulum (RyR2 and SERCA) Ca^2+^ contribution, compared to a 10%/90% ratio in mice. This makes rabbit Ca^2+^ handling less SR-dependent and more representative of human physiology. Epicardial adipose tissue is present in the human heart and scarce in mice. Rabbit hearts also have epicardial adipose tissue in the coronary groove, interventricular grooves and RV free wall.

**Figure S2.**
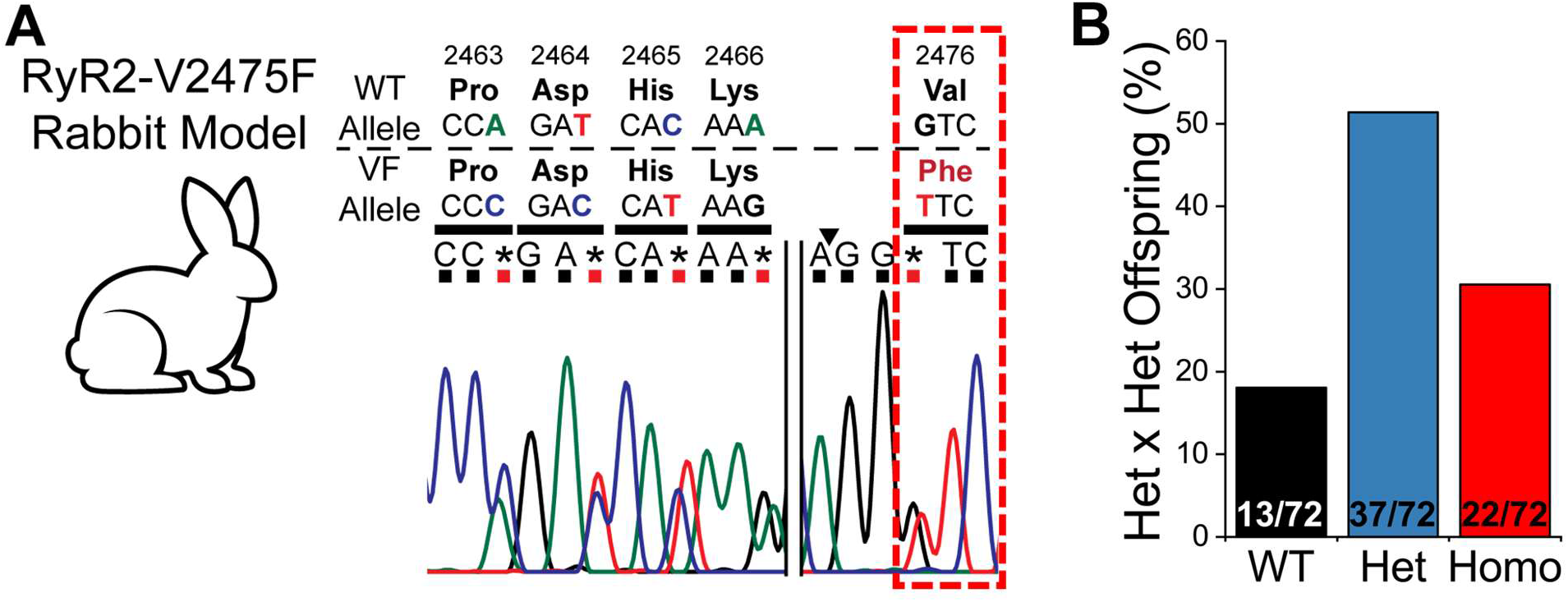
Generation of the RyR2-V2475F rabbit model. **A.** Strategy for model generation. RyR2-V2475F rabbits were created using CRISPR/Cas9 to introduce a missense mutation at codon 2476 (equivalent to human RyR2 codon 2475), resulting in a valine-to-phenylalanine (VF) substitution. This mutation abolished a Tth111I restriction site (arrowhead), which facilitated genotyping. Extensive genetic validation was performed to confirm the model. **B**. Offspring distribution of breeding heterozygote with heterozygote. N numbers as indicated in the column. Chi-square or Fisher Exact Test. ** P < 0.01. Het: heterozygote; Homo: homozygote.

**Figure S3.**
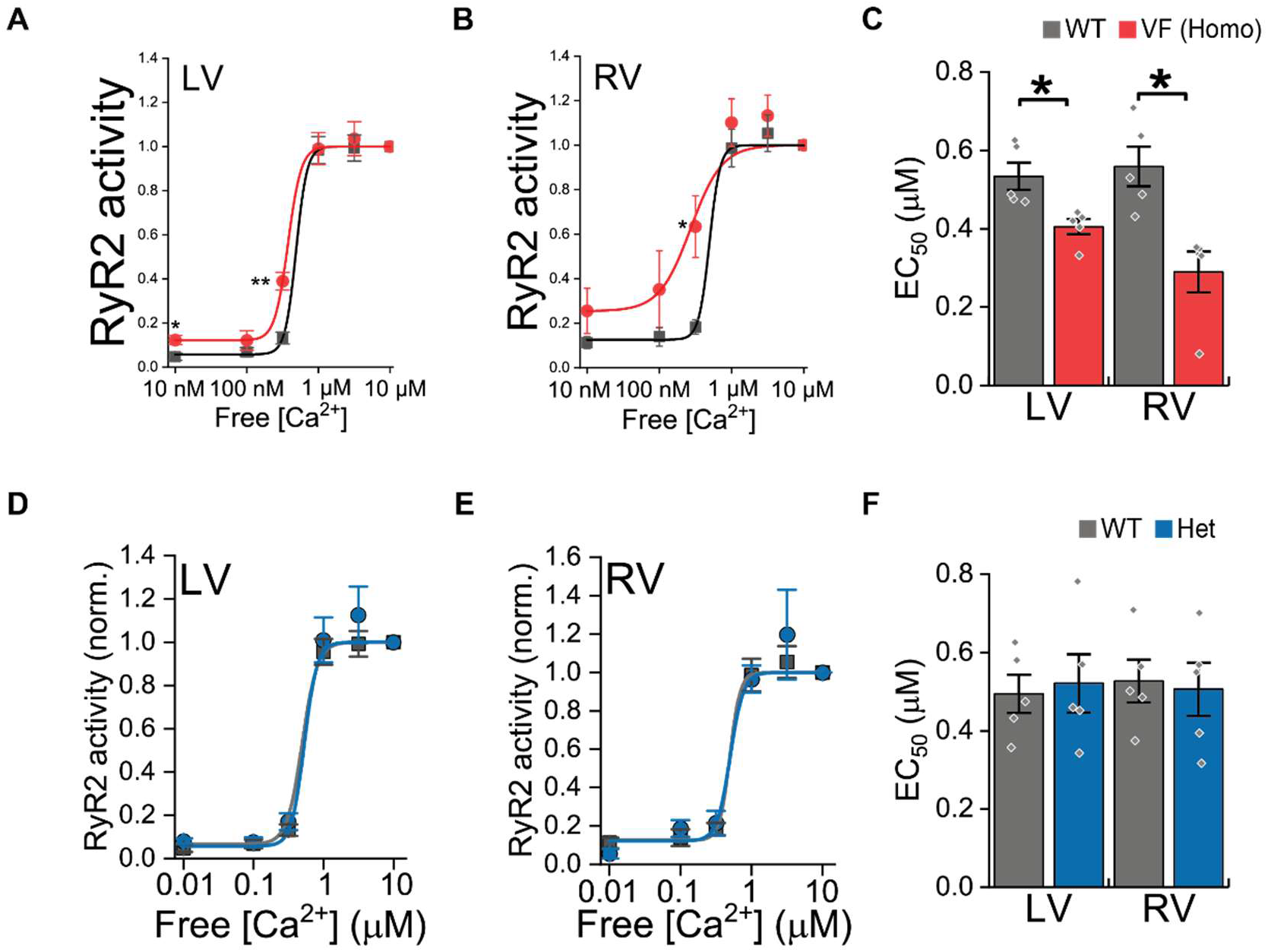
Enhanced RyR2 channel activity in V2475F knock-in (VF) homozygous hearts. **A-B**. RyR2 activity across a range of free [Ca^2+^], measured by [^3^H]ryanodine binding in LV (A) and RV (B) homogenates. Curves are normalized to maximal activation at 10 μM Ca^2+^. **C**. EC_50_ values for Ca^2+^-dependent RyR2 activation derived from sigmoidal fitting of curves in A-B. (mean±SEM; n = 5 per genotype. *P < 0.05; two-tailed t-test or Mann-Whitney test). D-F. [^3^H]ryanodine binding results for heterozygous (Het) hearts. n = 5 per genotype.

**Figure S4.**
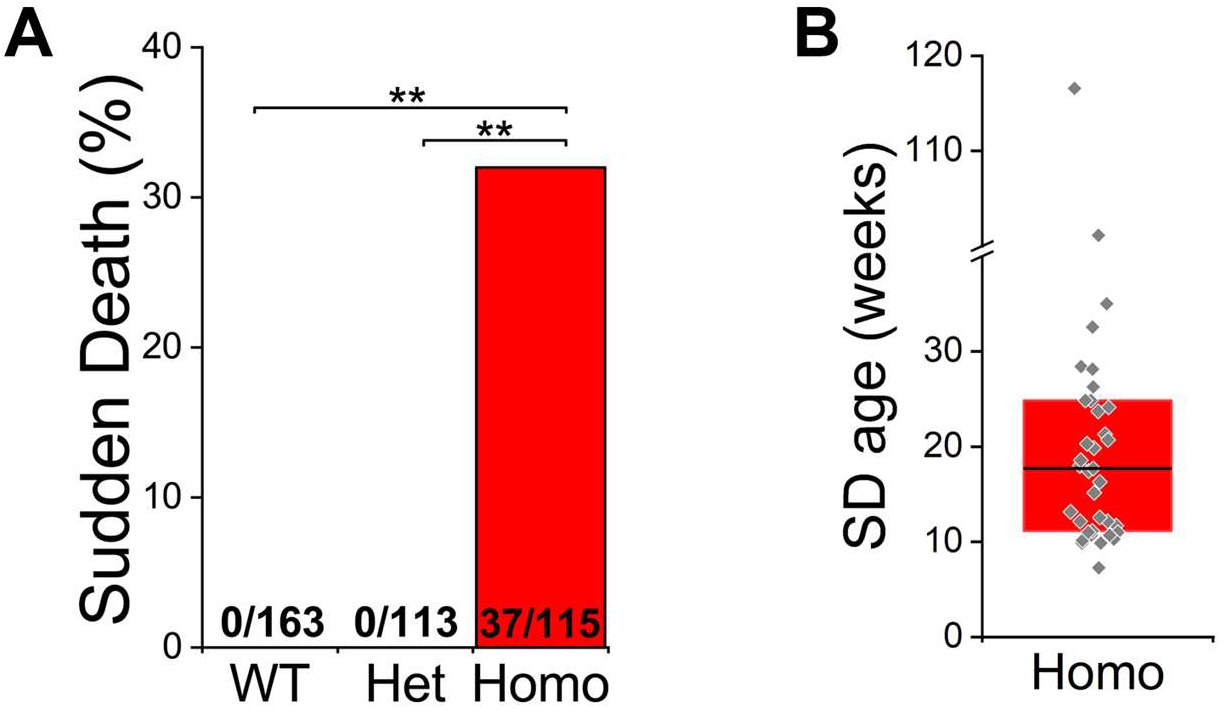
Characterization of sudden death (SD) in VF rabbits. **A.** Distribution of sudden death among all offsprings. N numbers as indicated in the column. Chi-square or Fisher Exact Test. ** P < 0.01. **B**. Age at sudden death in all RyR2-V2475F homozygous rabbits (N = 37). Ages ranged from 7.3 to 116.6 weeks, with a median age of 17.7 weeks. Het: heterozygote; Homo: homozygote.

**Figure S5.**
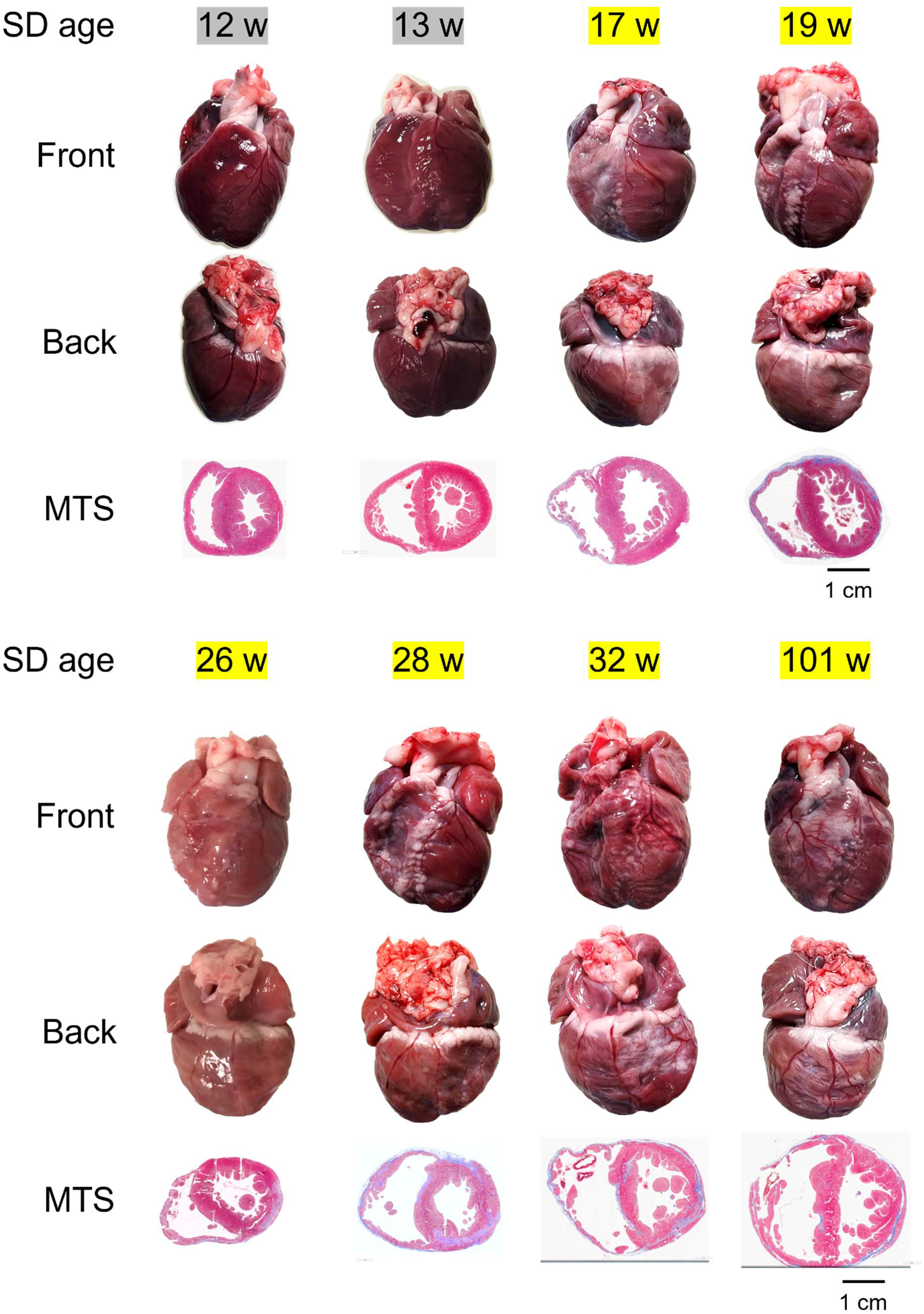
Hearts collected from RyR2-V2475F homozygous knock-in (VF)rabbits with sudden death (SD). **SD age:** Age at sudden death for each VF rabbit. Gray highlighting indicates the early stage; yellow highlighting indicates the advanced stage. **Front and Back:** Freshly extracted hearts from SD rabbits, showing front and back views. **MTS:** Masson’s trichrome staining (MTS) of transverse biventricular heart sections to visualize fibrosis. VF hearts exhibited progressive structural remodeling across stages. Early-stage (12–13 weeks, early) hearts showed RV dilation without fibrosis. With disease progression, hearts displayed RV-dominant wall thinning and fibrosis, eventually advancing to biventricular dilation with extensive fibrofatty infiltration. Muscle loss and infiltration followed an epicardium-to-endocardium gradient.

**Figure S6.**
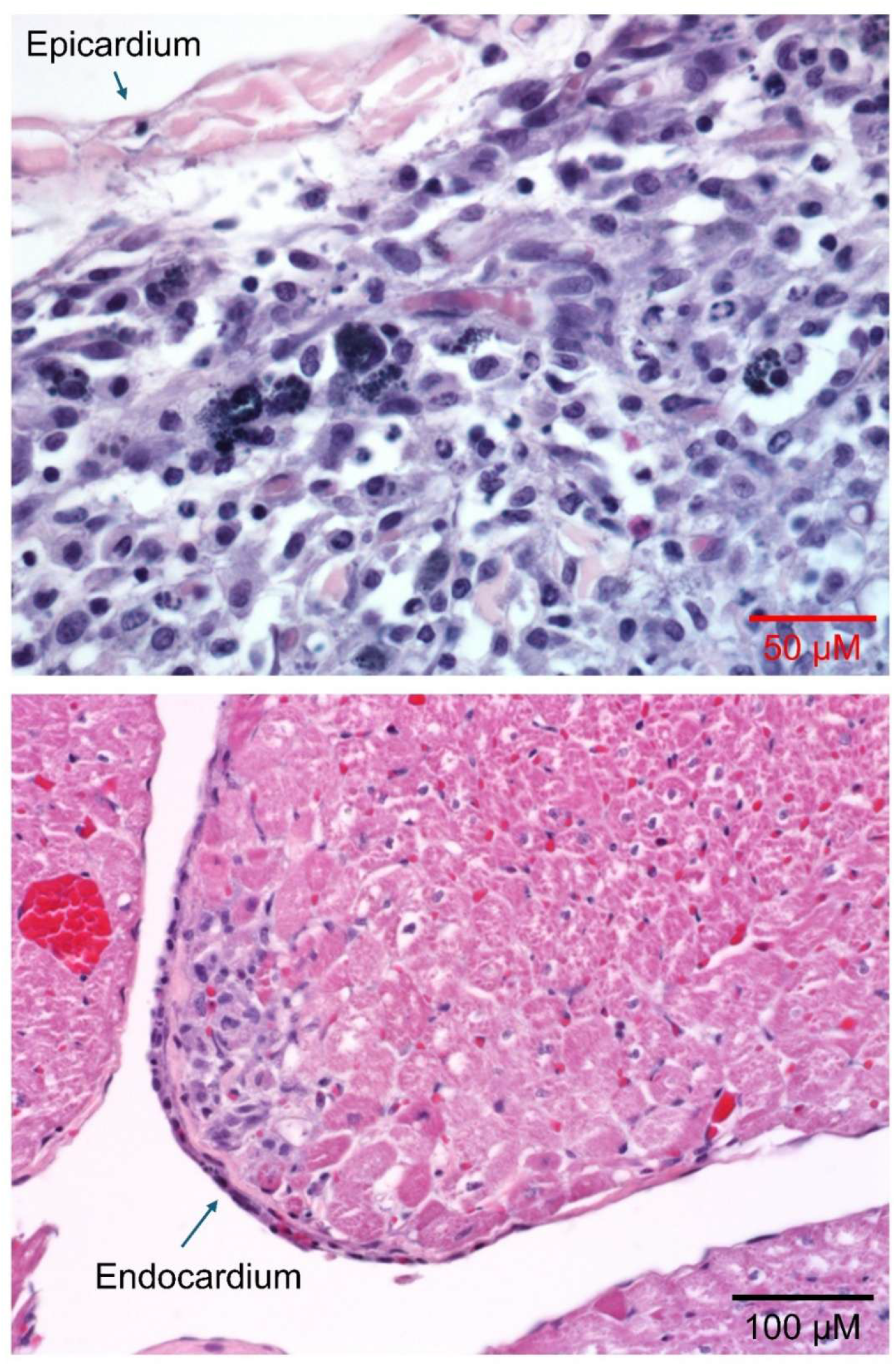
Patchy inflammatory infiltration in VF hearts from rabbits with sudden death. **Upper panel**: Monocyte infiltration observed in the epicardial region. **Lower panel**: Monocyte infiltration observed in the endocardial region. Slides are stained with Hematoxylin and Eosin (H&E).

**Figure S7.**
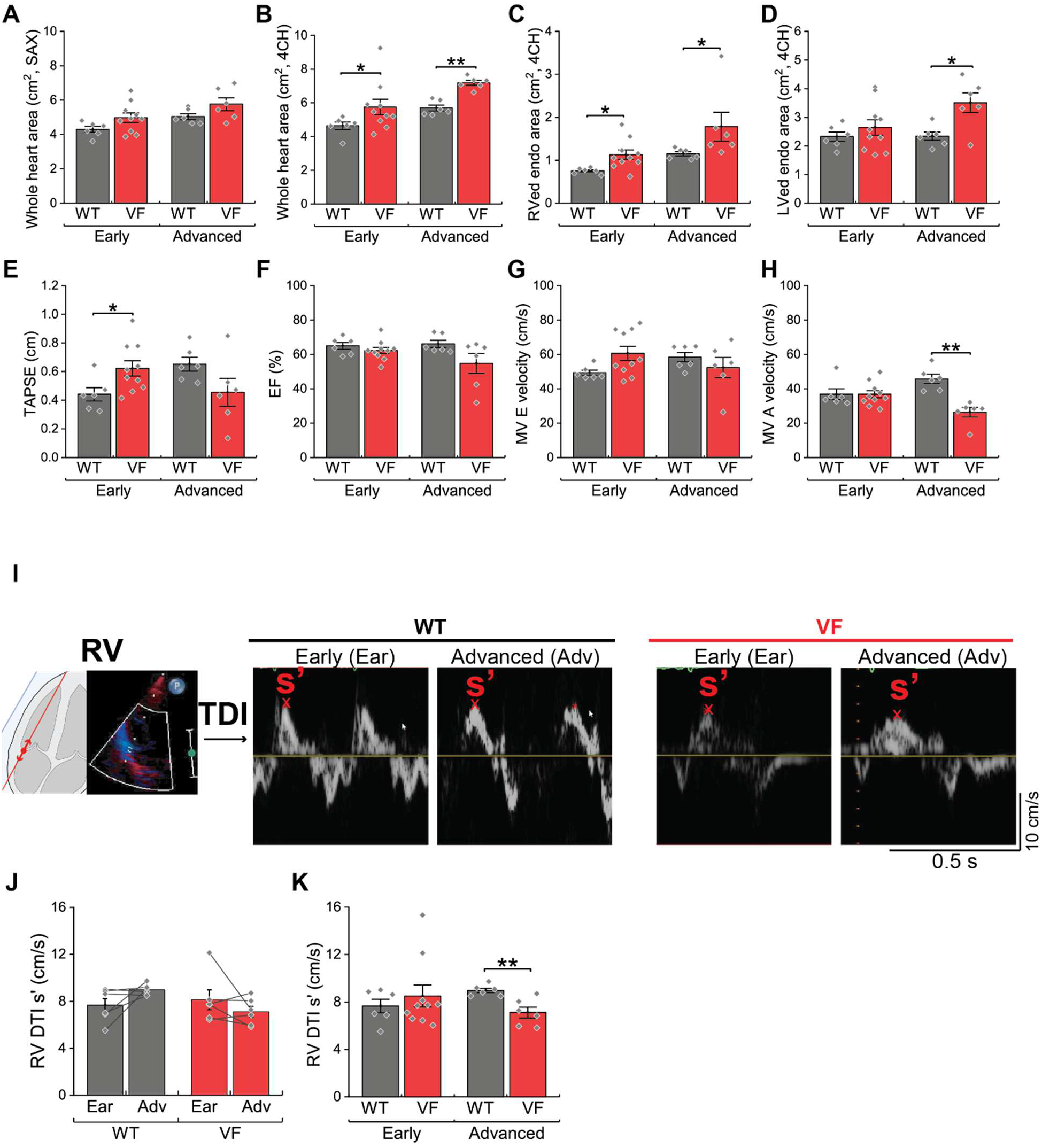
Cardiac echography measurements and RV tissue Doppler imaging (DTI) s′ velocity. **A-H.** Quantification of whole heart area (A-B), end-diastolic right ventricular endocardial (RVed endo) area (C), left ventricular endocardial (LVed endo) area (D), TAPSE (E), EF (F), MV E and A velocity (G-H). Comparisons are made between WT and VF in the matched stage. N = 6 WT (Early and Advanced); VF: 10 Early, 6 Advanced. **P < 0.05, *P < 0.01 by unpaired t-test or Mann-Whitney U test between genotypes. **I-K**. Representative echocardiographic imaging (I) and quantification (J-K) for RV DTI s’ velocity. RV DTI s′ velocity measures how fast the tricuspid valve moves during RV contraction, which was significantly reduced in VF rabbits at the advanced (Adv) stage. J: N = 6 each group. Paired T test or signed rank test within genotype. K: N = 6 WT (Early and Advanced); VF: 10 Early, 6 Advanced. Black asterisks indicate comparisons between age-matched WT and VF. *P < 0.05 by unpaired t-test or Mann-Whitney U test between genotypes.

**Figure S8.**
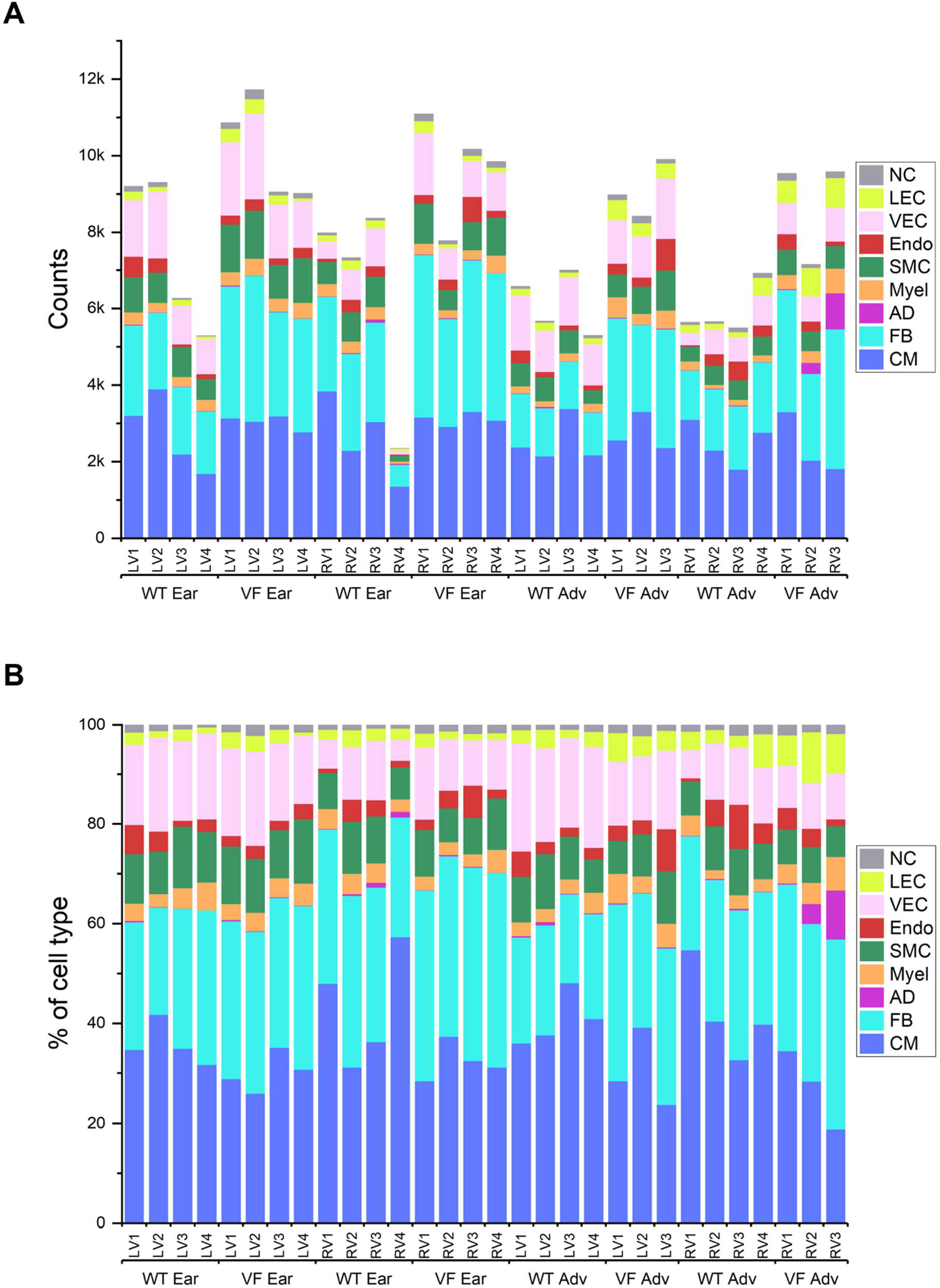
Nuclei count and cell type abundance for each sample. **A**. Nuclei counts from each cell type included for analysis in each sample after quality control and filtering. **B**. Cell type compositions for each sample, showing the relative abundance of each cell type. CM: cardiomyocytes, FB: fibroblasts, AD: adipocytes, Myel: myeloid cells, SMC: smooth muscle cells, Endo: endocardial cells, VEC: vascular endothelial cells, LEC: lymphatic endothelial cells, NC: neuronal cells.

**Figure S9.**
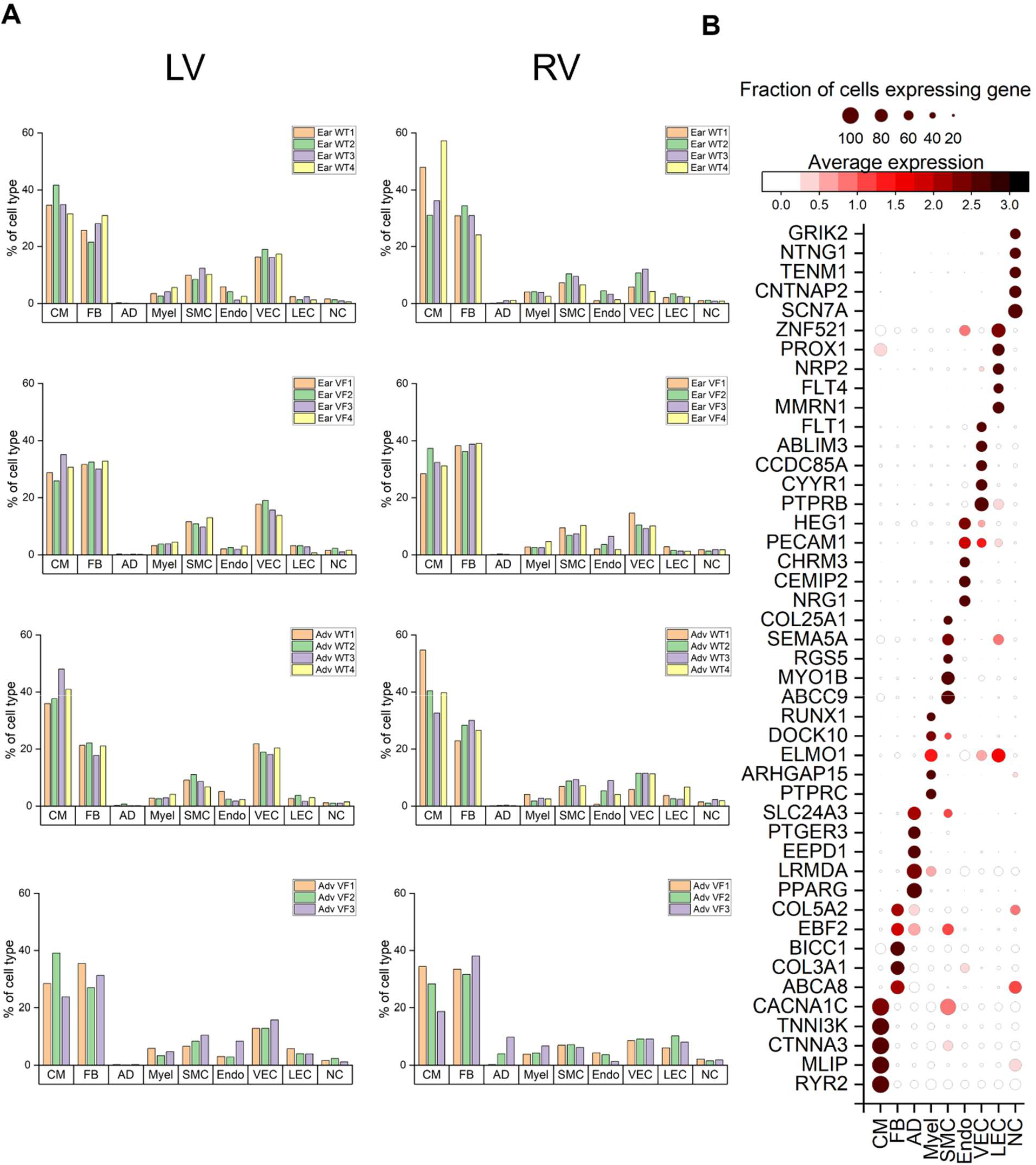
Cell type proportion of each sample and marker genes of each cell type. **A**. Cell type proportions for each sample from early (Ear) early WT and VF LV, Early WT and VF RV, advanced (Adv) WT and VF LV, Adv WT and VF RV. Groups are indicated in the plot. **B**. Dot plots showing selected marker genes for each cell type. CM: cardiomyocytes, FB: fibroblasts, AD: adipocytes, Myel: myeloid cells, SMC: smooth muscle cells, Endo: endocardial cells, VEC: vascular endothelial cells, LEC: lymphatic endothelial cells, NC: neuronal cells.

**Figure S10.**
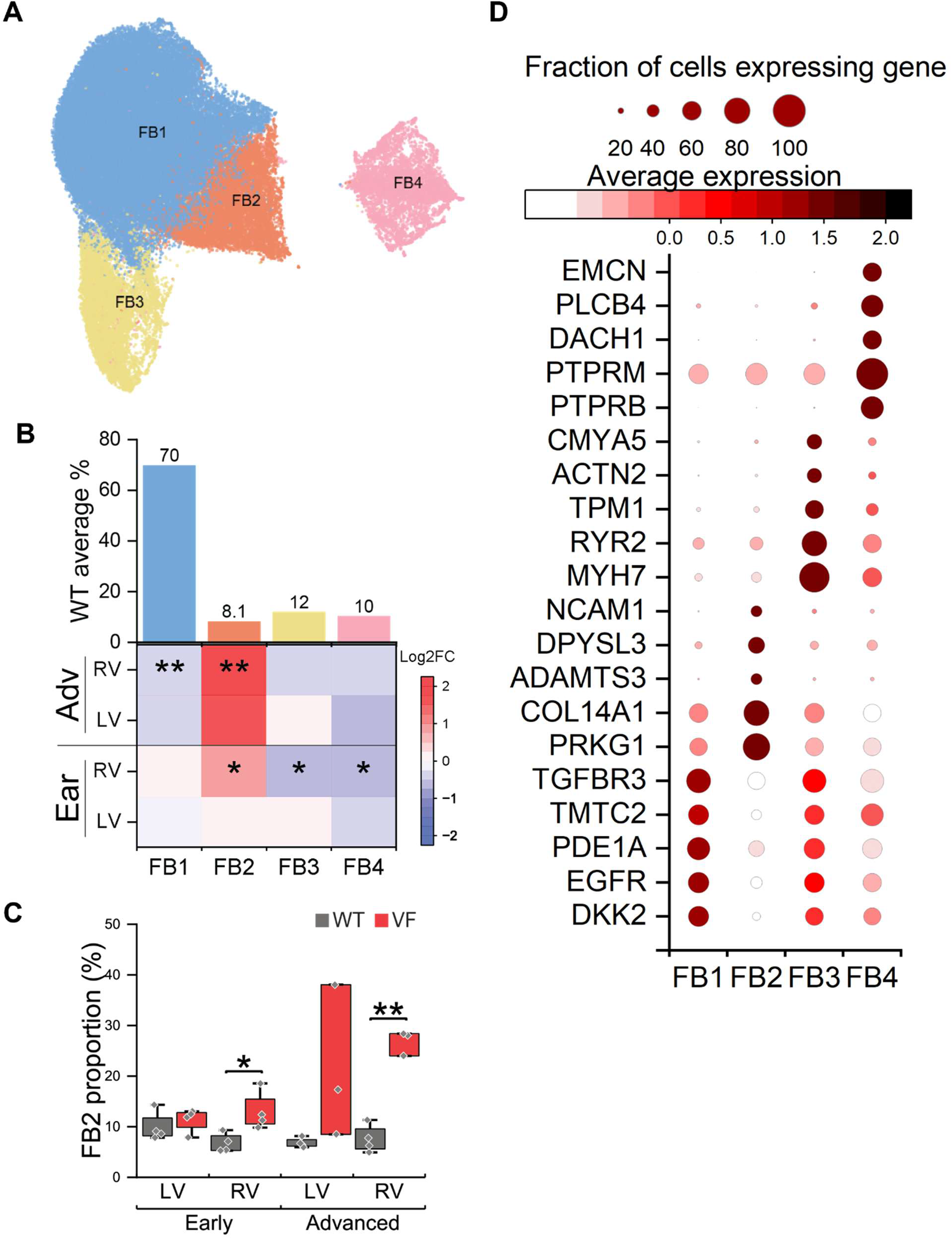
FB subclusters abundance and marker genes of each subcluster. **A-C**. Identification of fibroblast (FB) subclusters and stage-dependent abundance changes in snRNA-seq data. **A**: UMAP of FB subclusters (FB1-FB4). **B**: FB subcluster distribution in WT samples (upper panel) and heatmap of Log2(fold change, FC) changes in VF hearts compared to WT controls (lower panel; red, increased; blue, decreased). *P < 0.05, **P < 0.01 by T test or Mann-Whitney U test. **C**. Comparison of FB2 abundance between WT and VF. *P < 0.05, **P < 0.01 by T test or Mann-Whitney U test. **D**. Dot plots showing selected marker genes for each FB subcluster.

**Figure S11.**
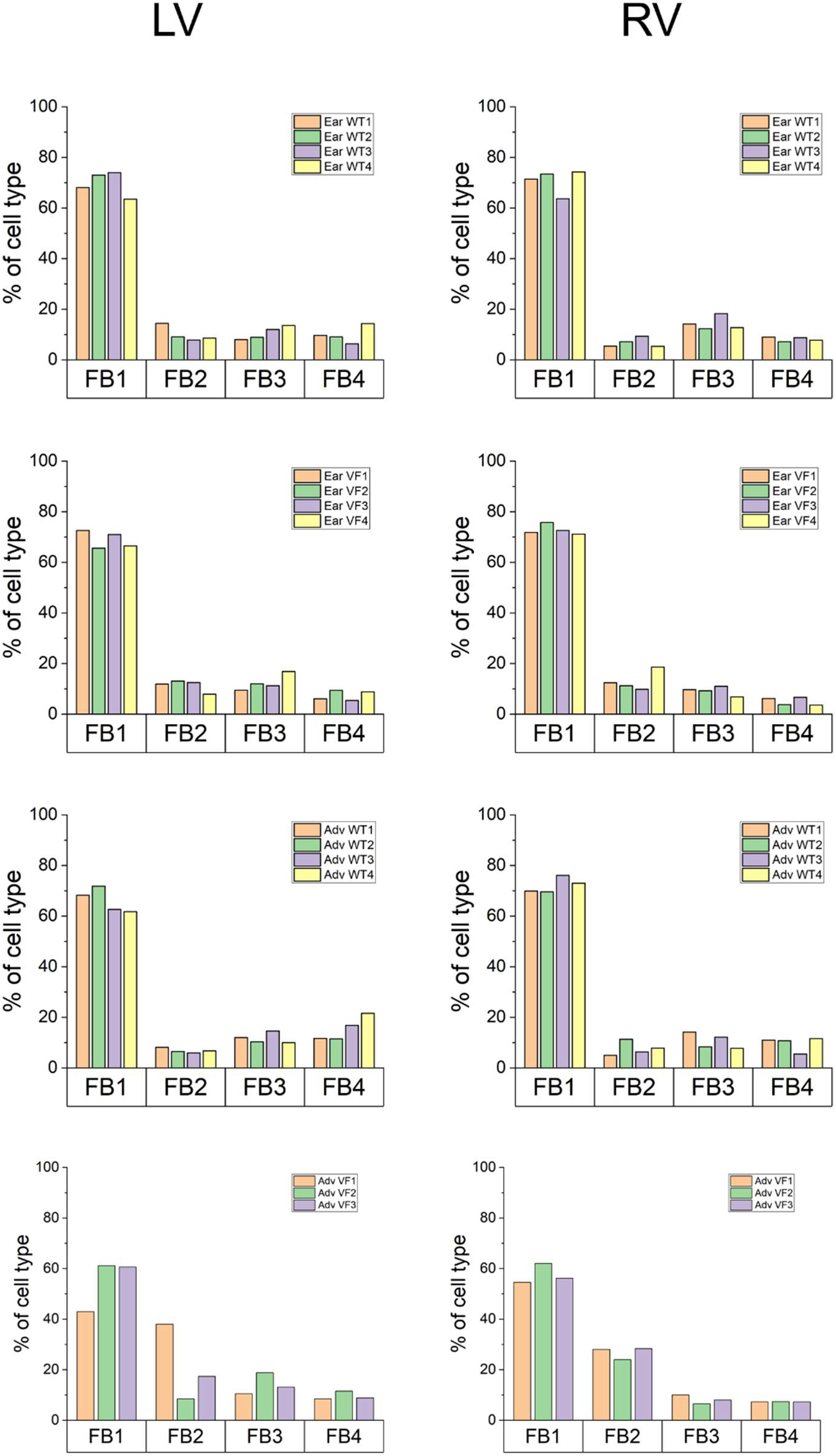
FB subcluster proportion of each sample. FB subcluster proportions for each sample from early (Ear) early WT and VF LV, Early WT and VF RV, advanced (Adv) WT and VF LV, Adv WT and VF RV. Groups are indicated in the plot.

**Figure S12.**
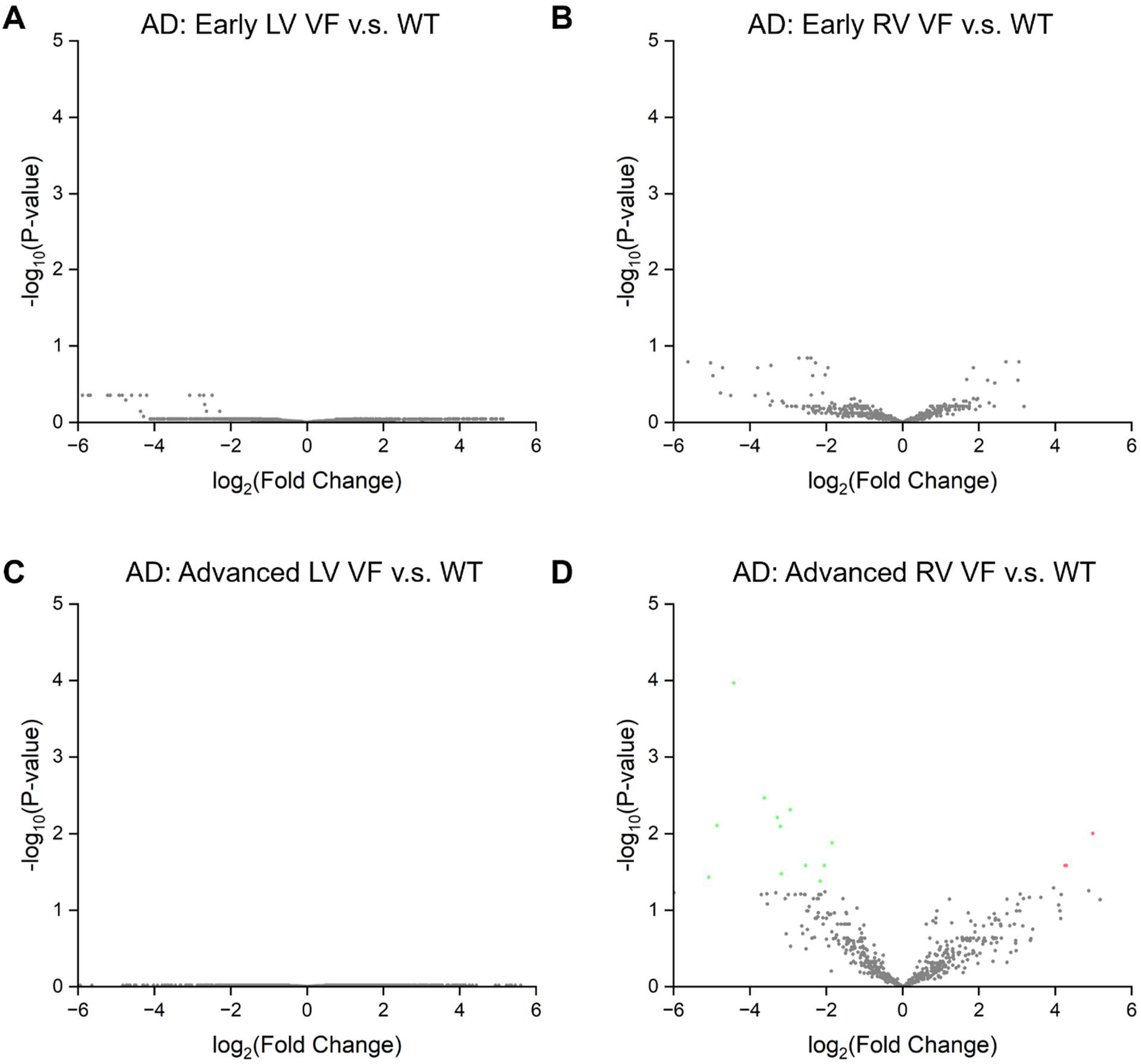
Volcano plot showing all and representative DEGs in Adipocytes (AD). **A-D**. Volcano plots showing all differentially expressed genes (DEGs) from early LV, early RV, advanced LV, and advanced RV. Not many DEGs were identified in adipocytes.

**Figure S13.**
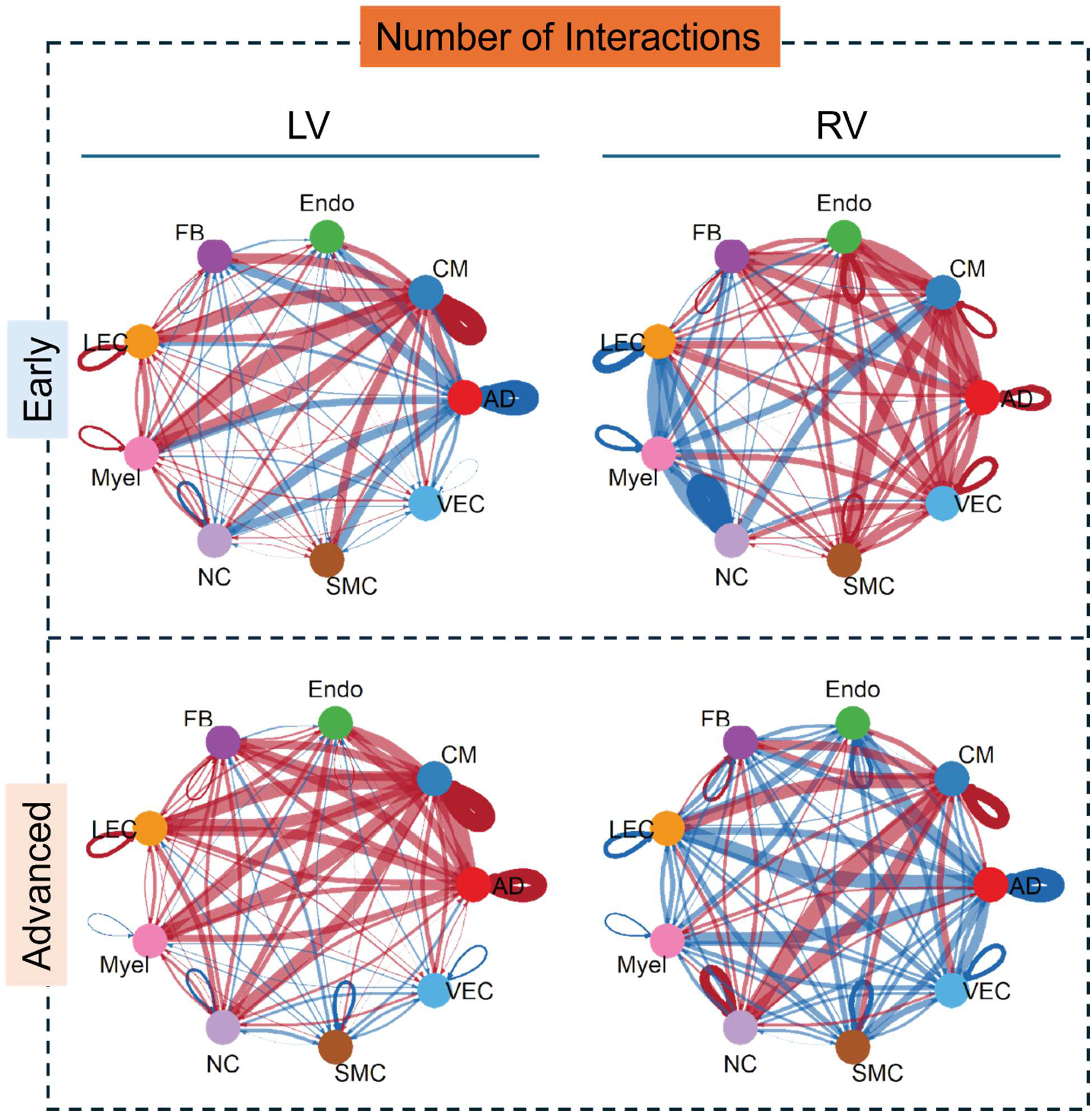
Differential interaction numbers between VF and matched WT by CellChat. Circle plots showing differential interaction numbers among major cell populations in VF versus matched control hearts. Red edges indicate increased signaling in VF; blue edges indicate decreased signaling in VF. Arrows denote directionality.

**Figure S14.**
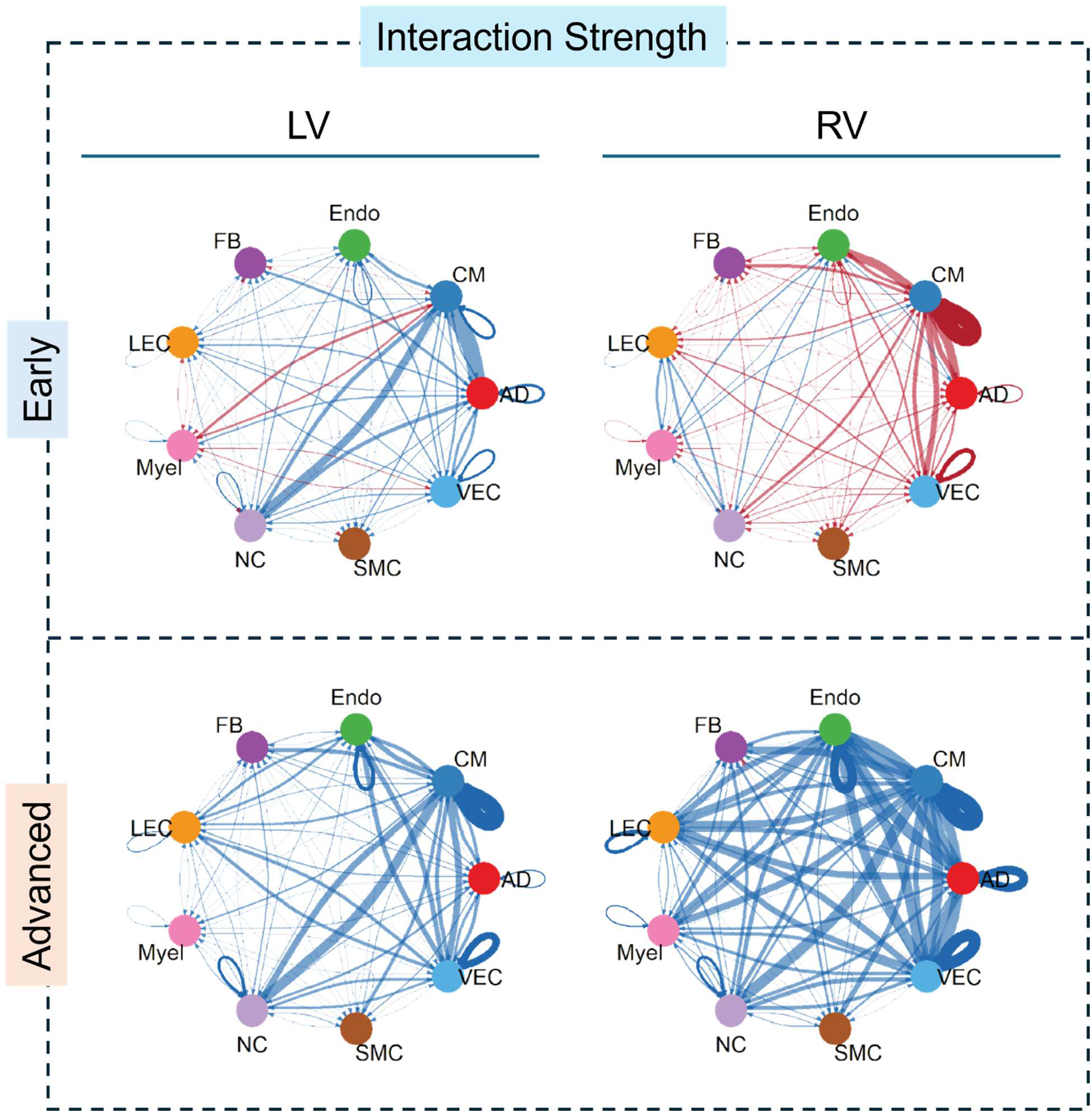
Differential interaction strength between VF and matched WT by CellChat. Circle plots showing differential interaction strength among major cell populations in VF versus matched control hearts. Red edges indicate increased signaling in VF; blue edges indicate decreased signaling in VF. Arrows denote directionality.

**Figure S15.**
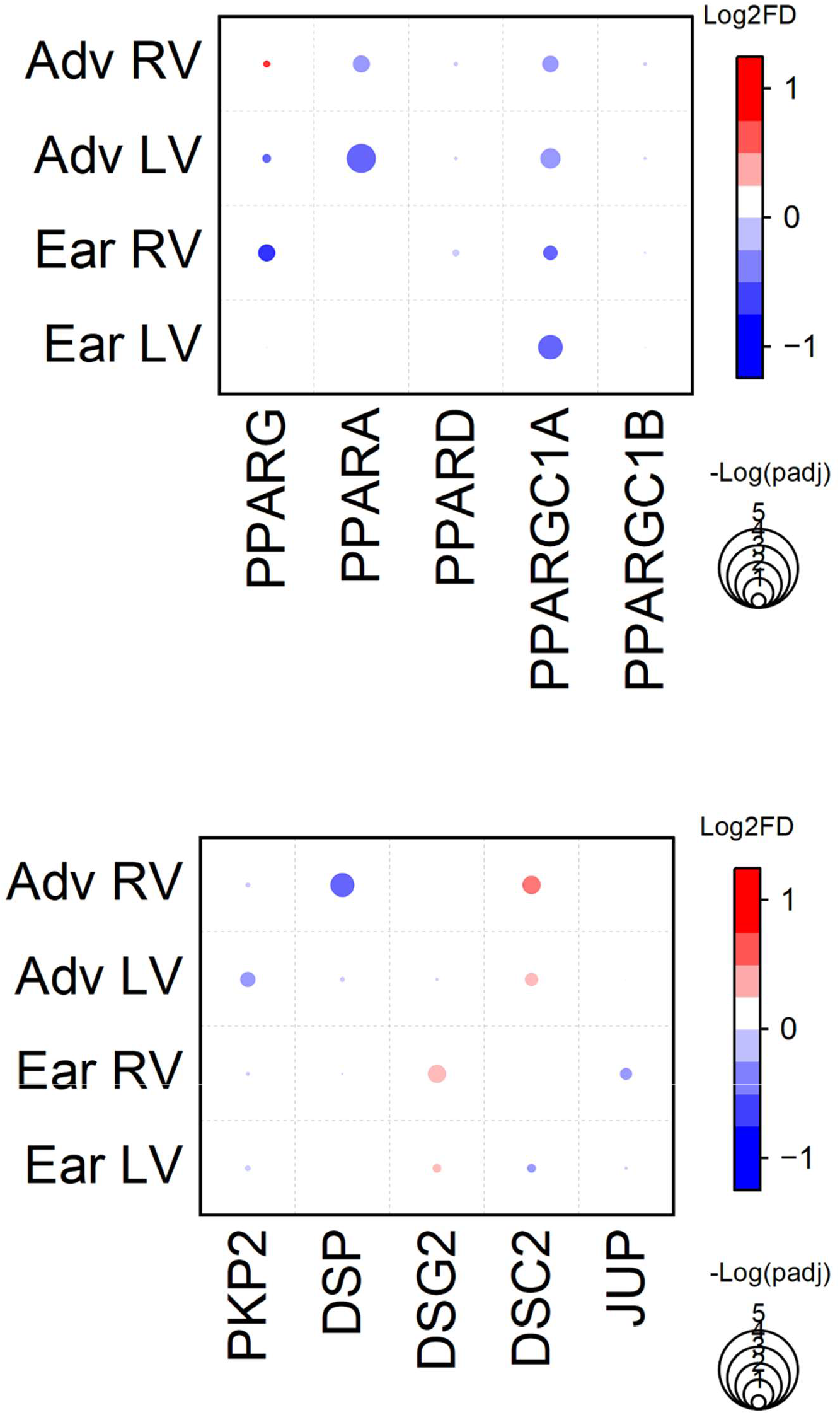
Bubble plots depicting expression changes of representative DEGs involved in PPAR signaling and major ACM pathogenic genes. Red indicates upregulation, while blue indicates downregulation. Bubble size indicates the -log10 adjusted p-value (-logPadj).

**Figure S16.**
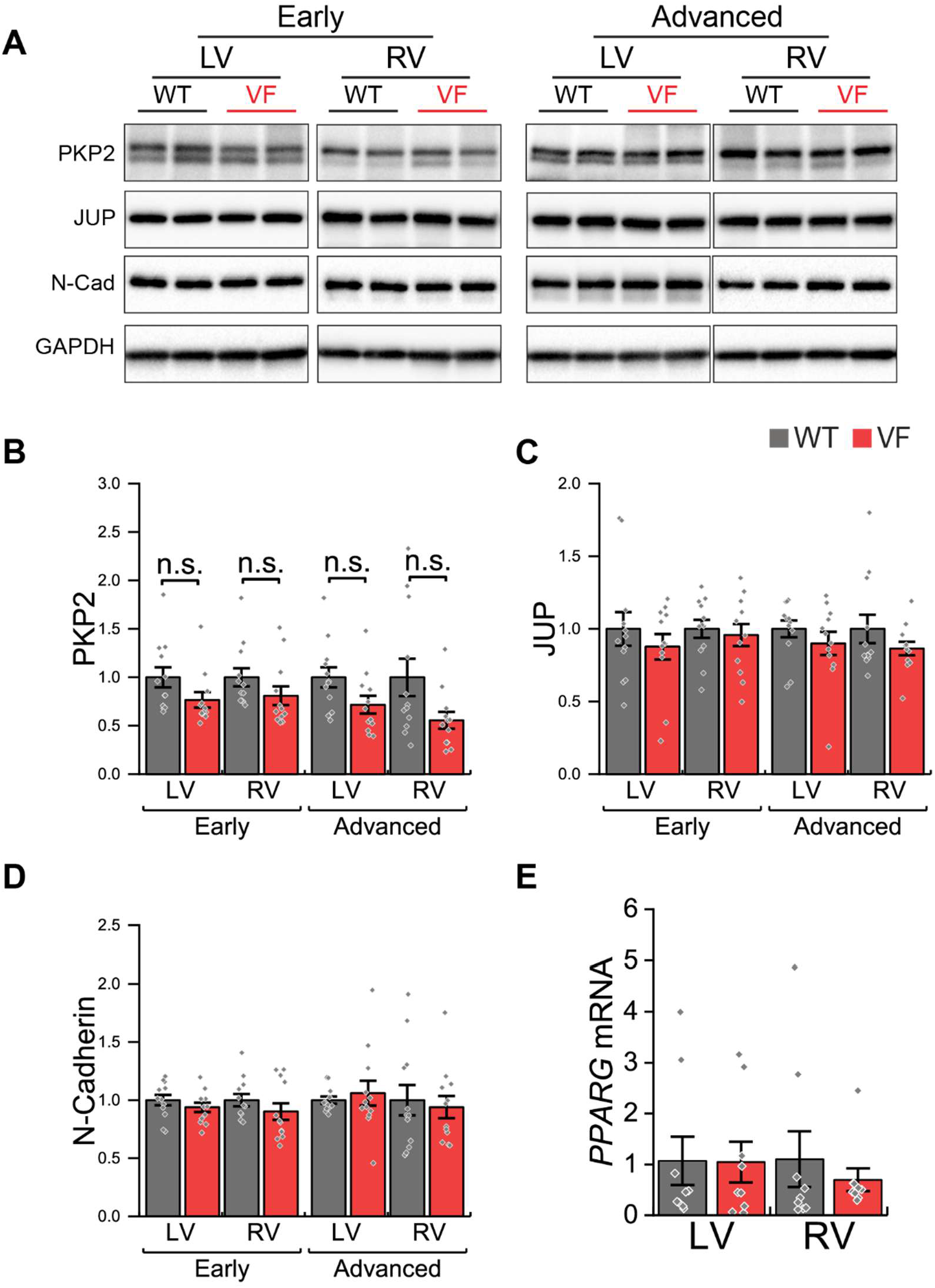
Desmosome and intercalated disc proteins expression. **A-D**. Representative immunoblots (A) and quantification (B-D) of intercalated disc (ICD) proteins. PKP2, plakophilin-2; JUP, junction plakoglobin; N-cad, N-cadherin. N = 12 per group. **E**. *PPARG* mRNA expression by qPCR from the early stage. N = 10 per group. Two-tailed t-test or Mann-Whitney test, n.s.: no significance.

**Figure S17.**
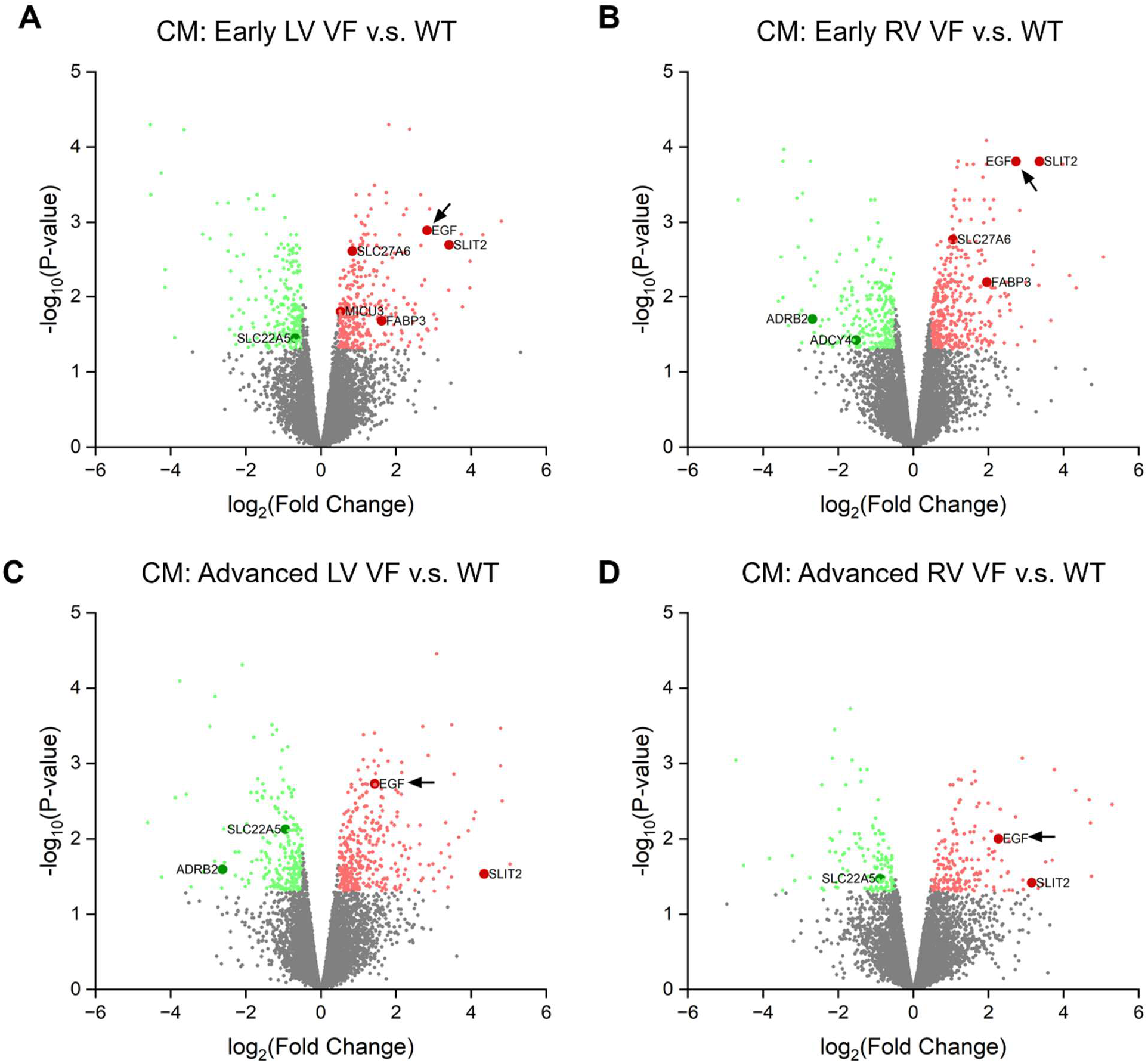
Volcano plot showing all and representative DEGs in CM. **A-D**. Volcano plots showing all differentially expressed genes (DEGs) in CM from early LV, early RV, advanced LV, and advanced RV. Significant DEGs are color-coded with red for upregulation and green for down regulation. Key genes involved in enriched pathways are highlighted and labeled with their gene names. Black arrow indicates EGF gene expression change.

**Figure S18.**
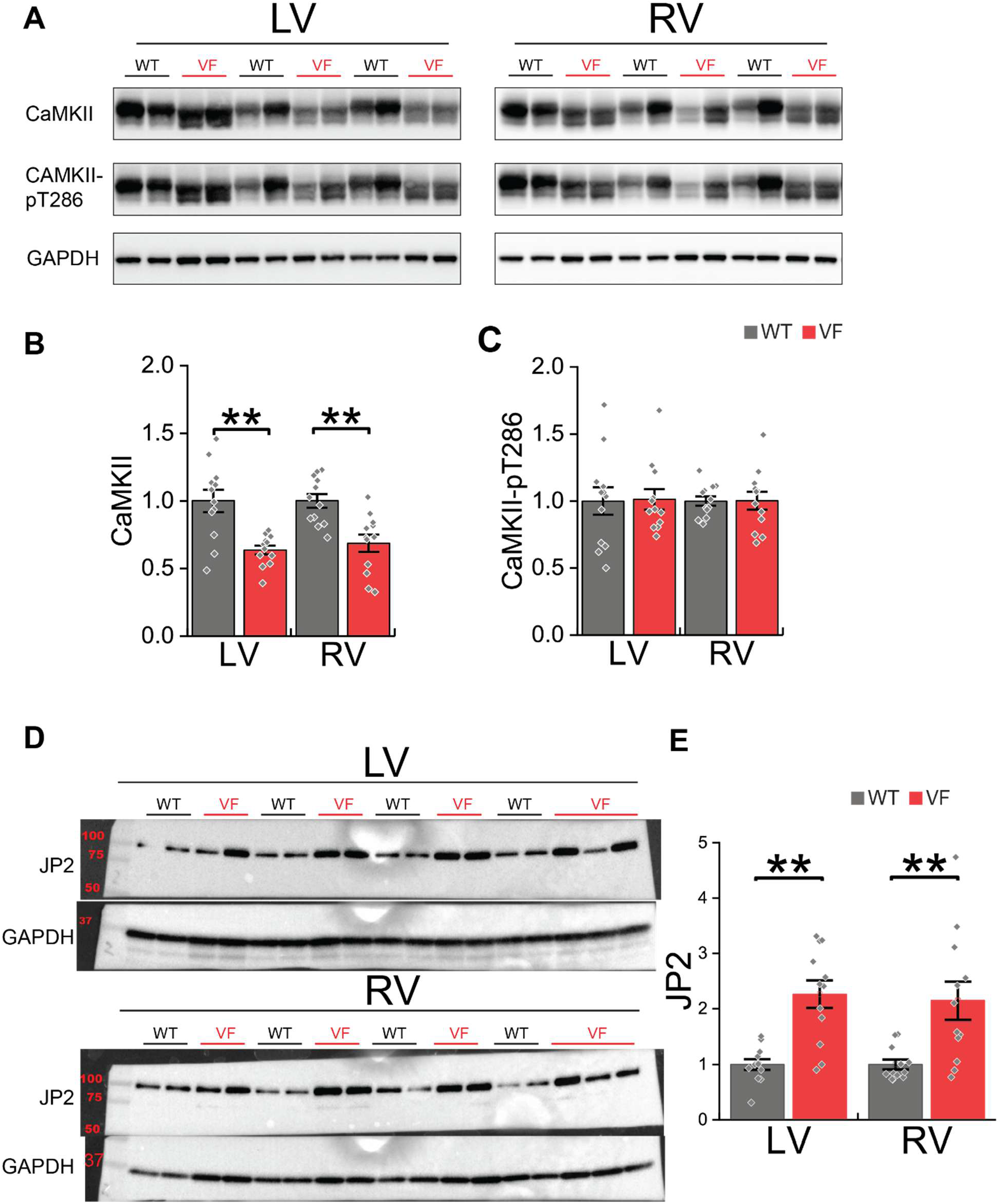
Marker proteins expression of canonical Ca^2+^-dependent stress in VF rabbit hearts. **A-C**. Representative Western blots and quantification of expression of CaMKII and autophosphorylated CaMKII at T286 (normalized to expression) in VF hearts at the early stage. Under pathological condition, Ca^2+^/calmodulin-dependent kinase II (CaMKII) activation is associated with increased cytosolic Ca^2+^ signaling. Yet, no evidence of increased total CaMKII or its activated form, as indicated by phosphorylation at T286. **D-E.** Representative Western blots and quantification of expression of Junctophilin-2 (JP2). JP2 and the cleaved product of N-terminal (JP2-NT) are well-established stress markers to Ca^2+^-dependent calpain activation. However, no sign of JP2 cleavage was observed, despite an increase in total JP2 protein. N = 12 per group. *P < 0.05, **P < 0.01; two-tailed t-test or Mann-Whitney test.

**Figure S19.**
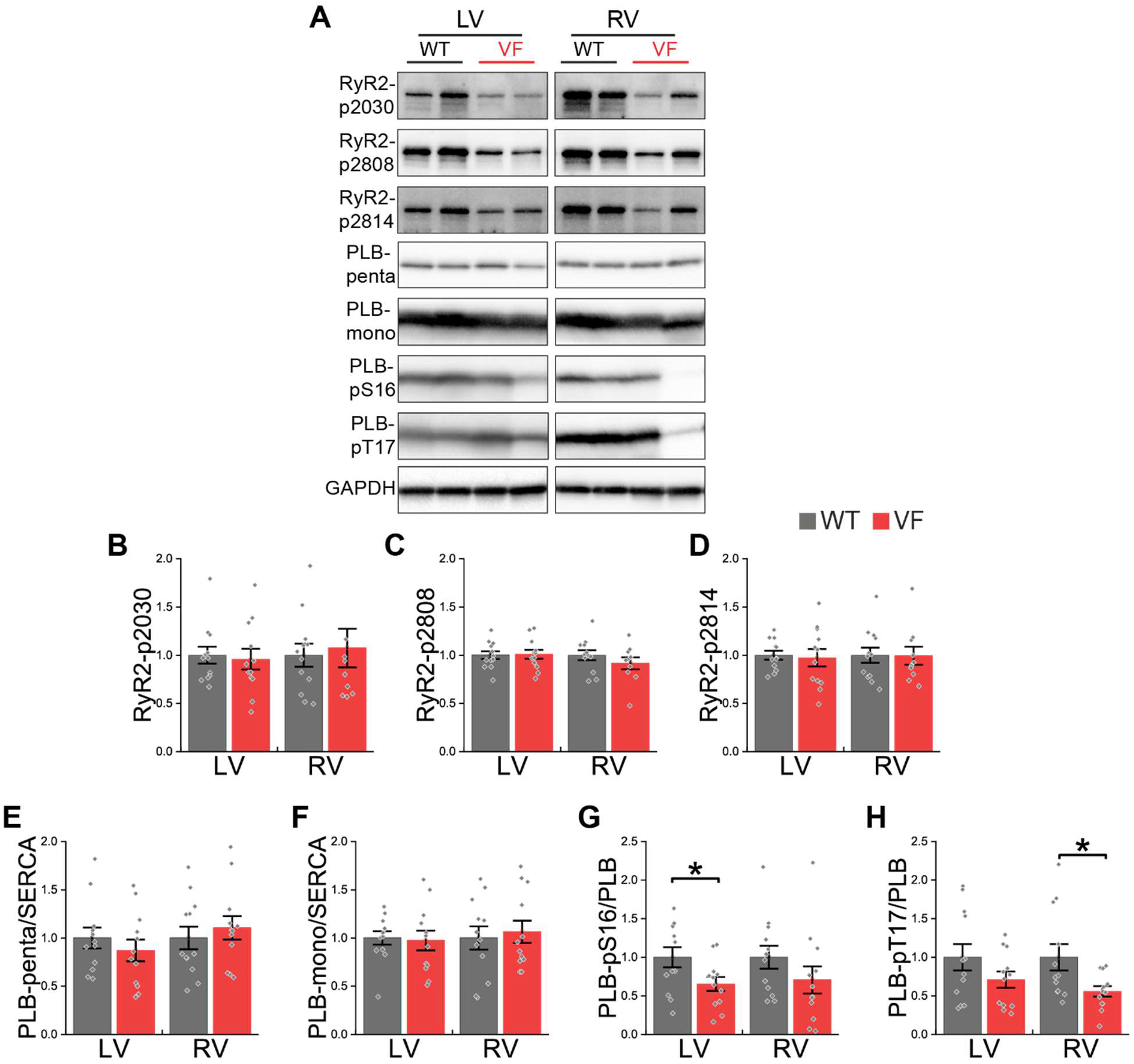
Phosphorylation levels of RyR2 and PLB. **A.** Representative Western blots of RyR2 phosphorylation, PLB expression (as pentamer or monomer) and PLB phosphorylation (only detectable in monomer). **B-H**. Quantification of expression and phosphorylation levels. Both RyR2 and PLB are well-established substrates of CaMKII. Yet, no increase in any of the well-known phosphorylation sites was observed on both proteins, further supporting the absence of CaMKII activation. N = 12 per group. *P < 0.05, **P < 0.01; two-tailed t-test or Mann-Whitney test.

**Figure S20.**
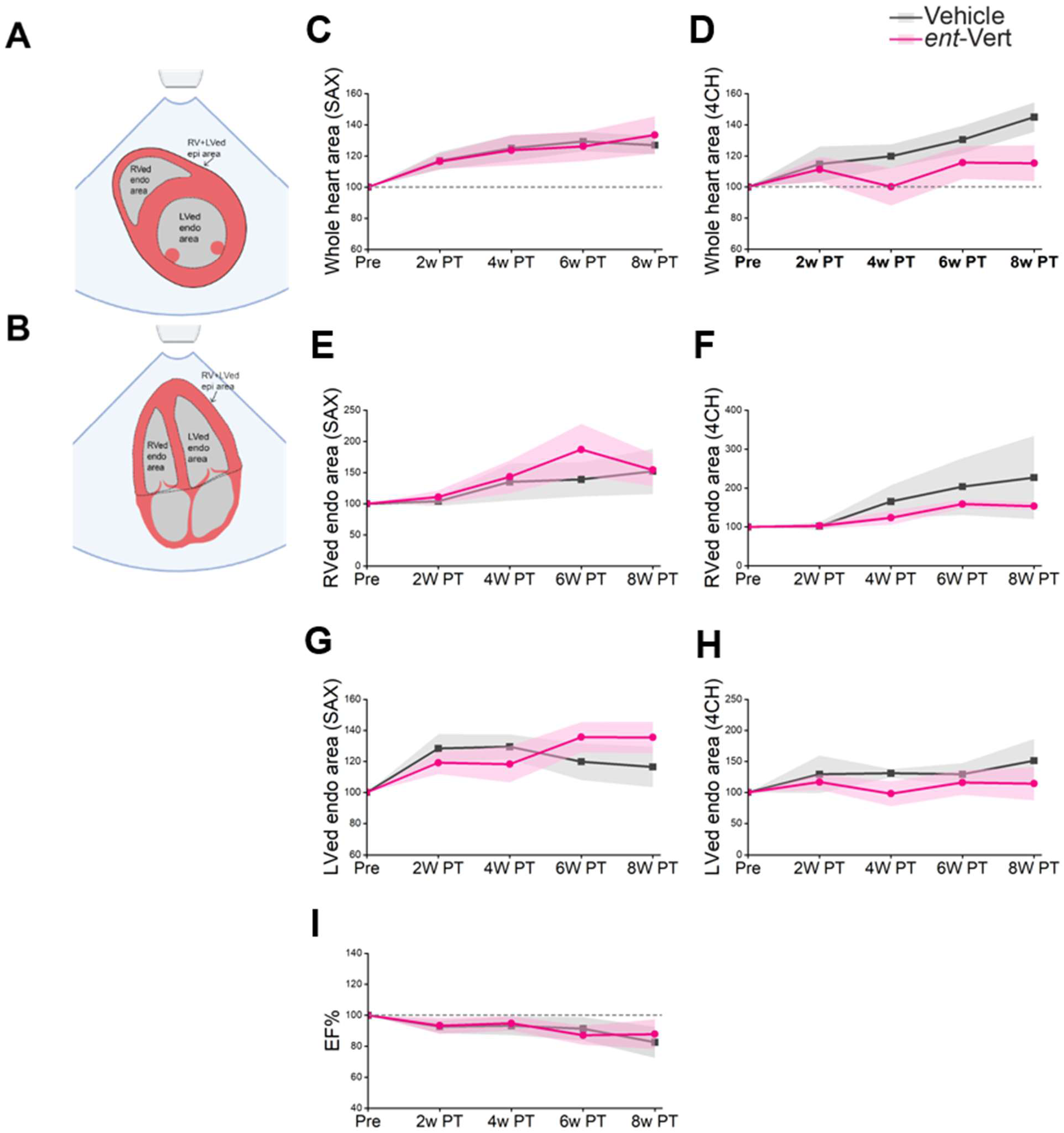
Progression of heart dimensions and EF by echocardiography during *ent*-verticilide treatment. **A–B.** Illustration of echocardiographic measurements of whole heart, left ventricular (LV) and right ventricular (RV) end-diastolic endocardial area in short-axis (SAX, A) and four-chamber (4CH, B) views. **C–H**. Longitudinal assessment of whole heart area (C, D) RV end-diastolic endocardial area (E, F) and LV end-diastolic endocardial area (G, H) over 8 weeks of treatment. Heart size and ventricular dimensions increased in both groups as rabbits grew during the monitoring period. No significant differences were observed between vehicle and treatment group. I. EF% progression. No significant differences were observed in EF between groups. Two way repeated ANOVA with mixed model. N = 5 per group. Shaded area around the lines indicates standard error.

**Figure S21.**
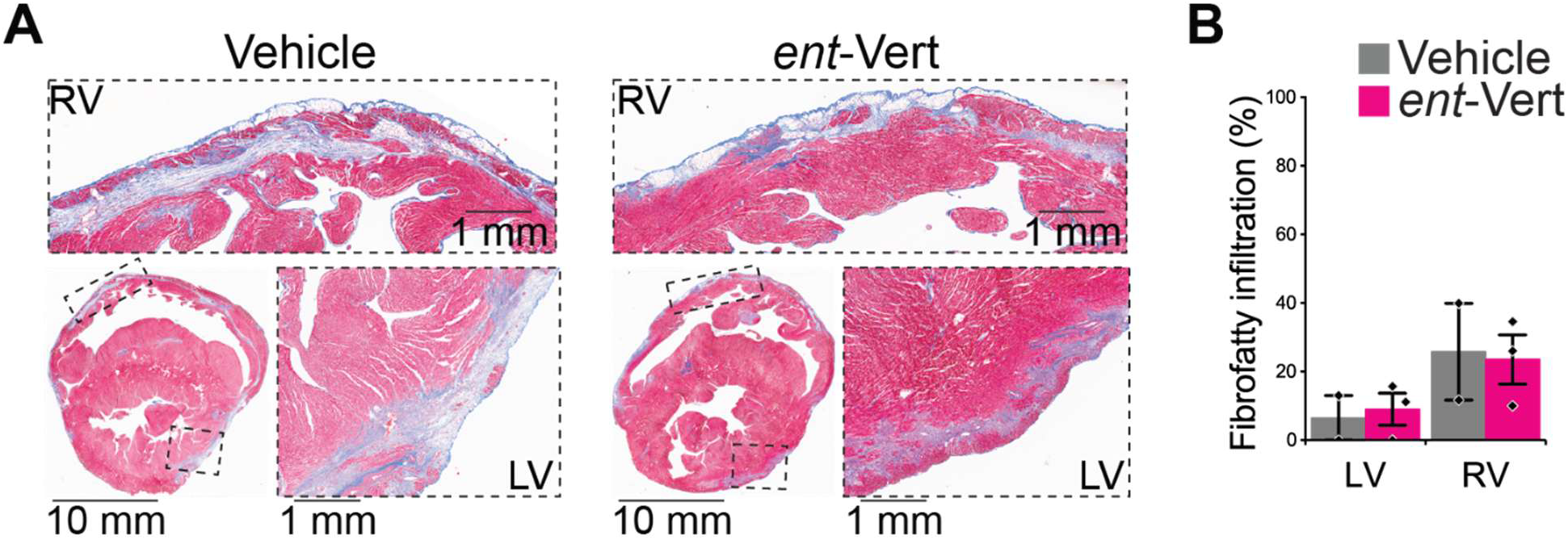
MTS staining and fibrofatty quantification from *ent*-verticilide treated rabbits. **A**. Representative Masson’s trichrome staining from vehicle and *ent*-verticilide treated hearts. Whole-heart sections and enlarged regions from RV (top) and LV (left) boxed areas are shown. B. Quantification of fibrofatty infiltration: present in all animals from both groups, with no significant difference. Vehicle N = 2 rabbits (1 from SD at the advanced stage, 1 from the surviving rabbit), *ent*-verticilide. N = 3 rabbits per group.

## Notes

### Competing Interest Statement

The authors have declared no competing interest.

